# A comparison of fecal glucocorticoid metabolite concentration and gut microbiota diversity in bonobos (*Pan paniscus*)

**DOI:** 10.1101/2022.01.04.474930

**Authors:** Alexana J. Hickmott, Klaree J. Boose, Monica L. Wakefield, Colin M. Brand, J. Josh Snodgrass, Nelson Ting, Frances J. White

## Abstract

Sex, age, diet, stress, and social environment have all been shown to influence the gut microbiota. In several mammals, including humans, increased stress is related to decreasing gut microbial diversity and may differentially impact specific taxa. Recent evidence from gorillas shows fecal glucocorticoid metabolite concentration (FGMC) did not significantly explain gut microbial diversity, but it was significantly associated with the abundance of the family Anaerolineaceae. These patterns have yet to be examined in other primates, like bonobos (*Pan paniscus)*. We compared FGMC to 16S rRNA amplicons for 201 bonobo fecal samples collected across five months to evaluate the impact of stress, measured with FGMC, on the gut microbiota. Alpha diversity measures (Chao’s and Shannon’s indexes) were not significantly related to FGMC. FGMC explained 0.08% of the variation in beta diversity for Jensen-Shannon and 1.27% for weighted UniFrac but was not significant for unweighted UniFrac. We found that genus *SHD-231*, a member of the family Anaerolinaceae had a significant positive relationship with FGMC. These results suggest that bonobos are relatively similar to gorillas in alpha diversity and family Anaerolinaceae responses to FGMC, but different from gorillas in beta diversity. Members of the family Anaerolinaceae may be differentially affected by FGMC across great apes. FGMC appears to be context dependent and may be species-specific for alpha and beta diversity but this study provides an example of consistent change in two African apes. Thus, the relationship between physiological stress and the gut microbiome may be difficult to predict, even among closely related species.

**Graphical Abstract:** See PDF

*Legend:* We compared the bonobo gut microbiota to fecal glucocorticoid metabolite concentrations (FGMC). FGMC did not explain alpha diversity, but FGMC explained ∼1.5% of the variation in beta diversity.

## Introduction

The gastrointestinal tract (gut) microbiota play essential roles in host nutrition and health across mammals [1]–[3]. Perturbations and dysbiosis to gut microbial communities have been linked to diseases like obesity, diabetes, irritable bowel disorders, and certain cancers in humans [4]–[8]. At the same time, physiological stress, hereafter referred to as stress, has been found to have a negative effect on the gastrointestinal tract and associated microbiota [9]. The definition of stress is a threat to homeostasis, which can be acute or chronic [10]. Chronic stress is stress that is experienced over a longer time frame (e.g., days to years), while acute stress often only lasts for several minutes to hours [11], [12]. As used in the human and non-human primate (NHP) literature, stress often refers to psychosocial stress or stress caused by a response to social stimuli that disrupts the normal physiological equilibrium. We will be using stress to refer to chronic stress, including psychosocial stress and other types of stress like dietary stress. However, determining the cause of stress can be difficult, and a very fine level of both behavioral and biomarker data is needed to determine the source of stress. Therefore, we will be only referring to chronic stress broadly when saying stress. Stress has been linked to dysbiosis, or a disruption to the homeostasis of the microbial community, in the gut microbiota in several mammalian taxa, including humans [13]–[17], and has also been implicated in mediating the communication between a host and their commensal gut microbiota [18], [19]. The gastrointestinal tract and its associated microbiota play essential roles in host nutrition and health across mammals [1]–[3].

Perturbations and dysbiosis to gut microbial communities have been linked to diseases like obesity, diabetes, irritable bowel disorders, and certain cancers in humans [4]–[8]. At the same time, stress has been found to have a negative effect on the gastrointestinal tract and associated microbiota for humans in the clinical setting [9]. Stress hormones have been proposed as a mechanism for communication along the gut-brain axis [15]. The gut-brain axis is a bidirectional signaling pathway between the brain and the gut that is potentially mediated by gut microbiota [20]. For example, in humans, stress has been linked to a decrease in the number of species found in the gut microbiota [9]. Additionally, evidence in humans suggests a link between stress, the gut microbiota, and immune system function [21]. This evidence from humans is also recapitulated in laboratory models [11], [22]–[25]. In germ-free rats, the lack of a gut microbial community increases a rat’s behavioral and endocrine stress response [22]. Other lab studies in mice have linked depression symptoms, anorexia, and cancer to increased stress levels and disruptions in the murine gut microbiota [24], [26], [27].

Stress and the gut microbiota has primarily been examined in laboratory models, with very few studies looking at wild living mammals [15], [25]. In wild-living eastern grey squirrels, physiological indicators of stress were measured in conjunction with gut microbial composition, and physiological stress better explained gut microbiota diversity, or metrics that summarize how abundant and the types of bacterial and archaeal species that are in a microbial community, compared to environmental factors [28], [29]. A recent study on pangolins found those raised in captivity had higher microbial diversity associated with lower stress than those rescued from the wildlife trade [30]. In elephants, a stressful event such as translocation appears to induce alterations to the microbiome in taxa Planococcacea. Clostridiaceae, Spirochaetaceae, and Bacteroidia increased after the elephants shifted to living in a captive environment [31]. Research into glucocorticoid concentration in rhinos found that about 10% of taxa were related to glucocorticoid concentrations with Aerococcaceae, Atopostipes, Carnobacteriaceae, and Solobacterium differentially increased [32]. Therefore, there seems to be a general mammalian pattern that suggests higher stress is associated with increases in specific taxa. However, this pattern has not been well-studied across mammals, and the specific taxa that increase seem to vary widely across mammalian groups.

Amongst NHPs, great apes exhibit similar psychosocial and ecological pressures making them an excellent model to understand how stress and the gut microbiome co-vary [33]–[35]. The mammalian pattern of higher stress being associated with a more disrupted gut microbiome [28], [30], [31], [36] was not found in a study of wild-living western lowland gorillas (*Gorilla gorilla gorilla*) [10]. Vlčková et al. (2018) found no relationship between alpha and beta diversity measures and proximate stress measures in this species. Vlčková and colleagues also found a positive correlation between proximate measures of stress and relative abundance of three different gut microbial taxa (family Anaerolineaceae, genus *Clostridium* cluster XIVb and genus *Oscillibacter*), suggesting that stress is associated with increases in certain types of bacteria within the gastrointestinal tract [10]. These results from western lowland gorillas indicate stress may not have as significant an effect on NHP gut microbial diversity but follows the mammalian trend of having specific taxa differentially abundant with increasing stress. Bonobos (*Pan paniscus*) and gorillas face similar social stressors in that they both are group-living great apes experiencing affiliative and aggressive interactions with conspecifics [37]. They also tend to live in mixed age and mixed-sex groups, though in western lowland gorillas, there is typically only one male [38]–[43]. Whether all NHPs have the same taxa that increase with increasing stress and whether they show similar stability in how the diversity of the gut microbiota remains stable has yet to be determined. Additionally, the amounts and effects of stress hormones vary across different groups of mammals and NHPs and may be different in closely related taxa, like gorillas and bonobos.

Stress hormones, specifically glucocorticoids such as cortisol, are considered the principal chemical compounds involved in the stress system of mammals, including NHPs [18], [44]. Glucocorticoids are a type of steroid hormone that fall into the class of corticosteroids, and cortisol, the primary mammalian stress hormone, is one type of glucocorticoid involved in the stress response [18]. Once the adrenal gland excretes cortisol, it is broken down and metabolized [45]. These metabolites are then excreted through saliva, feces, and urine and can be incorporated into tissues like nails and hair [46]–[49]. These metabolites in feces are referred to as FGMC, and are known to be related to chronic stress rather than acute stress [50], [51]. Therefore, FGMC are capturing the stress an individual experienced in the preceding forty-eight hours before a fecal sample was collected [52]. It is of note that sometimes low stress can be an indicator of suppression of the stress response as well as low stress. The relationship between the gut microbiota and hormonal systems has far-reaching implications for host physiology [53]. Nevertheless, how a host’s gut microbiota responds to various stress-based fluctuations during short-term variation in stress remains to be examined in many NHPs, like bonobos.

Stress has been studied in wild-living bonobos as it relates to sociality and socio-sexual behavior [33], [46], [54], [55]. Bonobos are female bonded, male philopatric, and exhibit a fission-fusion social system, and this social structure may contribute to the sex-based patterns in FGMC [39], [40]. Sex differences in bonobo stress have been quantified in inter-group encounters with both females and males exhibiting higher cortisol during intergroup encounters but with males having overall higher levels of cortisol [56]. Additionally, captive bonobos exhibited a similar pattern where the single male had overall higher cortisol levels than the five other females [57]. These physiological patterns and social structure more likely emulate that of the *Pan*-*Homo* common ancestor as compared to gorillas making bonobos an ideal model in which to study stress and the microbiome.

We aim to use bonobos as a model to test patterns of NHP stress and gut microbiota. We predict that bonobo gut microbiota will exhibit a pattern similar to what was found in western lowland gorillas because gorillas and bonobos live in a similar social and ecological environment. Gorillas and bonobos have been hypothesized to use similar resources, like terrestrial herbaceous vegetation, perhaps buffering dietary stress for both great ape populations, unlike chimpanzees [58], [59]. Bonobos and gorillas may have similar gut microbiota because they are both great ape species or because of similar environments. Bonobos and gorillas diverged 8-19 million years ago and, therefore, may exhibit differences due to phylogenetic differences [60], [61]. However, the gut microbiota may alternatively have similar responses to stress due to these ecological similarities between bonobos and gorillas. If they are very different, then it suggests that ecological environments are not the important factor in a gut microbiota’s response to FGMC. We predict that alpha diversity, or within individual diversity, will not be significantly related to FGMC. We predict that beta diversity, or between individual diversity, will not be significantly related to FGMC. We expect to find several taxa’s abundance, specifically family Anaerolineaceae, genus *Clostridium* cluster XIVb and genus *Oscillibacter,* to be significantly explained by FGMC.

## Methods

### Study Site and Sample Collection

The research site was the Iyema field camp, located in the Lomako-Yokokala Faunal Reserve just north of the Lomako river at (00°55) North, 21°06) East) in the Democratic Republic of Congo (DRC). The site was mainly covered by primary forest in *terra firma* soil with swamps [55], [62].[55], [62]. We followed bonobos to their night nests for data collection as part of the African Wildlife Foundation’s habituation efforts from June 2017 to October 2017. Night nest locations were marked, and each nesting site was revisited the following day. We identified each bonobo as it exited the nest and collected approximately five grams of fecal sample into 50 mL tubes with 10 ml of RNALater for each individual in the nesting party [63]. While there is some debate about the effectiveness of different sample preservation methods for examining gut microbiomes [62]-[68], there is no clear present consensus. We used RNAlater here due to field site remoteness and downstream host genetic analyses.

The samples were stored in a cool, dry place from June-October 2017 until shipped to the Ting Laboratory at the University of Oregon. They were then stored in a minus 20°C freezer until extraction. The remainder of each fecal sample not collected in RNALater was brought back to camp, dried using a camp stove, and placed into bags with desiccant for FGMC analysis. Thus, for each fecal sample, we can obtain data on gut microbiota composition and diversity and FGMC. We collected 218 paired fecal samples.

### Data Collection

#### ELISA assays - FGMC

To evaluate FGMC, we analyzed 218 dried fecal samples in the Global Health Biomarker Laboratory at the University of Oregon using ELISA assays to quantify cortisol as a measure of FGMC. We used the Arbor Assay’s DetectX^®^ Cortisol Enzyme Immunoassay Kit (Arbor Assay’s DetectX^®^ cat. no. K003-H5W), as it is designed to be used on dried fecal samples and was previously validated for bonobos [71], [72]. We included known controls provided for Cincinnati Zoo bonobos (N=5) for each plate run. Fecal samples were ground to a powder using a mortar and pestle, weighed out to the protocol’s recommended ≥ 0.2 g. of fecal material, avoiding any plant or partially digested food material. Samples were then diluted (1:4) in assay buffer. The kit manufacturer reported the detection limit for this assay as 45.4 pg/mL. To control for shifts in circadian rhythm for FGMC, we used those samples collected at the same time of day, specifically in the morning, under night nests, to ensure all bonobo samples were from approximately the same time point. All plates were read using a BioTek microplate reader and analyzed with Gen5 software version 2.0. For the FGMC controls, 100□μl aliquots of assay buffer were divided into seven aliquots. We then spiked six of the aliquots with 100□μl of standards such that each aliquot of the sample received one of the six concentrations of standard (1000, 500, 250, 125, 62.5, 31.2□pg/mL), and one aliquot was left neat following the kit protocol to produce a standard curve. Both the spiked and neat aliquots were assayed according to kit instructions.

### 16S sequencing – gut microbiota composition

We used the 218 RNALater preserved fecal samples to extract, amplify, and sequence microbial DNA. Total genomic DNA was extracted from each fecal sample using the QIAamp PowerFecal DNA kit (QIAGEN) in the Ting Lab at the University of Oregon. Negative controls were included in extraction batches to test for contamination. DNA extracts were quantified using a Qubit dsDNA HS Assay Kit protocol using a Qubit 2.0 Fluorometer (Thermo Fisher Scientific). Samples containing at least 1.0 ng/μl were sent for amplification and sequencing of the V4 hypervariable region of the bacterial 16S ribosomal RNA using 515F/806R primers at the Genomics and Cell Characterization Core Facility at the University of Oregon following previously published methods [73], [74]. Barcoded amplicons were sequenced on a 150 PE V3 run on the Illumina MiSeq platform (Illumina, San Diego, CA). The resulting sequences were demultiplexed and denoised using DADA2, and amplicon sequence variants (ASVs) were assigned using the Green Genes database. Quality filtering and assembly were done using the QIIME2 pipeline for microbial analyses [75, p. 2]. The ASVs table was created for samples rarified to an average sampling depth of 79,058 reads per sample. We removed samples below 315 and above 100,000 reads per sample which accounted for 8.7% of the original dataset. We retained 202 samples for a total of 26,010,213 reads. Negative control samples were sequenced for each extraction, PCR, and library preparation. Any ASVs that appeared in these negative controls were removed from the 202 samples in the R package ‘decontam’ using the prevalence and frequency methods [76].

### Data analysis

We tested sex, age, whether a female had an infant, and fecal glucocorticoid metabolite concentration (FGMC) as our predictor variables with bonobo gut microbiota composition and diversity as the response variable. We included these variables because they are associated with differences in FGMC and gut microbiota [77]–[81]. Statistics were run in R version 4.0.2 [82]. We included sample ID as a random effect in our models [83]. We estimate 26-38 individuals sampled with an estimated resampling rate of 2-11 times during the data collection period based on observations of individuals and previously published estimates for Iyema [84]. We calculated two measures of alpha diversity, Shannon’s and Chao’s diversity indices, using the ‘vegan’ package [85]. We ran two-way ANOVAs against the predictor variables, against sex (N = 55) and age (N=59), sex (N = 55) and whether a female had an infant (N=59), and age (N=59) and whether a female had an infant (N=59). We ran linear regressions for FGMC (N=202) against Shannon’s index and Chao’s index to study alpha diversity or within individual diversity (Table 1). To examine the relationship between FGMC and bonobo gut microbiota beta diversity, we ran a permutational multivariate analysis of variance (PERMANOVA) with 999 permutations using the ‘adonis2’ function in the R package [85]. PERMANOVAs use the calculated beta diversity for the Jensen-Shannon Distance, weighted UniFrac, and unweighted UniFrac dissimilarity matrices taking the model predictors FGMC, sex, age, and whether or not a female had an infant sequentially (n=202) (Table 1). We used Jensen-Shannon’s Distance because it is useful for examining compositional differences [86], [87]. Weighted UniFrac is a metric used to detect differences based on commonly abundant taxa. At the same time, unweighted UniFrac is better at detecting differences in rare or non-abundant taxa in a community. [88]. It is of note that PERMANOVAs factor in the order in which variables are entered into the model, therefore we ran this with the factors in different orders and found that the pattern of significance stayed consistent despite the order of the predictor variables. We also ran Mantel tests on the log transformed FGMC values and the three beta dissimilarity matrices. We also ran abundance models on our filtered data, and ran 302 taxa using the ANCOM R package to test whether a member of the gut microbiota varies with high (21,540 – 7,115 ng/pl and low concentrations (1,073 – 7,115 ng/pl) of FGMC (N=202) [55], [89]. The cut-off was used because ANCOM requires a categorical variable of about equal sample sizes to run. All models were run with FGMC as a continuous variable except the ANCOM results which required the FGMC values to be coded as high or low. We then subsetted out the significant ASVs and verified with general linear model that those significant taxa had a linear relationship with FGMC.

**Table 1.** Predictor variables included in analysis and collection method. Samples sizes are in parentheses.

| Predictor variable | Factor levels | Collection method | Lab analysis |
| --- | --- | --- | --- |
| FGMC | Continuous value (202) | Non-invasive fecal sample collection | ELISA assays to quantify cortisol |
| Sex | Male (24), female (31) | Behavioral observations corroborated with genetic sexing assay | Sexing assay |
| Age | Adult (48), sub-adult (4), juvenile (5), infant (2) | Behavioral observations | -- |
| Infant (Whether or not a female had an infant) | Yes (10), no (49) | Behavioral observations | -- |

## Results

### Alpha diversity

Bonobo’s fecal microbiome had similar alpha diversity regardless of FGMC levels, their sex, age or whether a female had an infant. Shannon’s and Chao’s indexes were not significantly related to FGMC (Figure 1). The ANOVA results were all not significant for Shannon’s and Chao’s diversity indices against sex (Shannon: p-value = 0.919; Chao’s: p-value = 0.488), age (Shannon: p-value = 0.955; Chao’s: p-value = 0.699), and whether or not a female had an infant (Shannon: p-value = 0.912; Chao’s: p-value = 0.521) (Figure 2).

**Figure 1.**
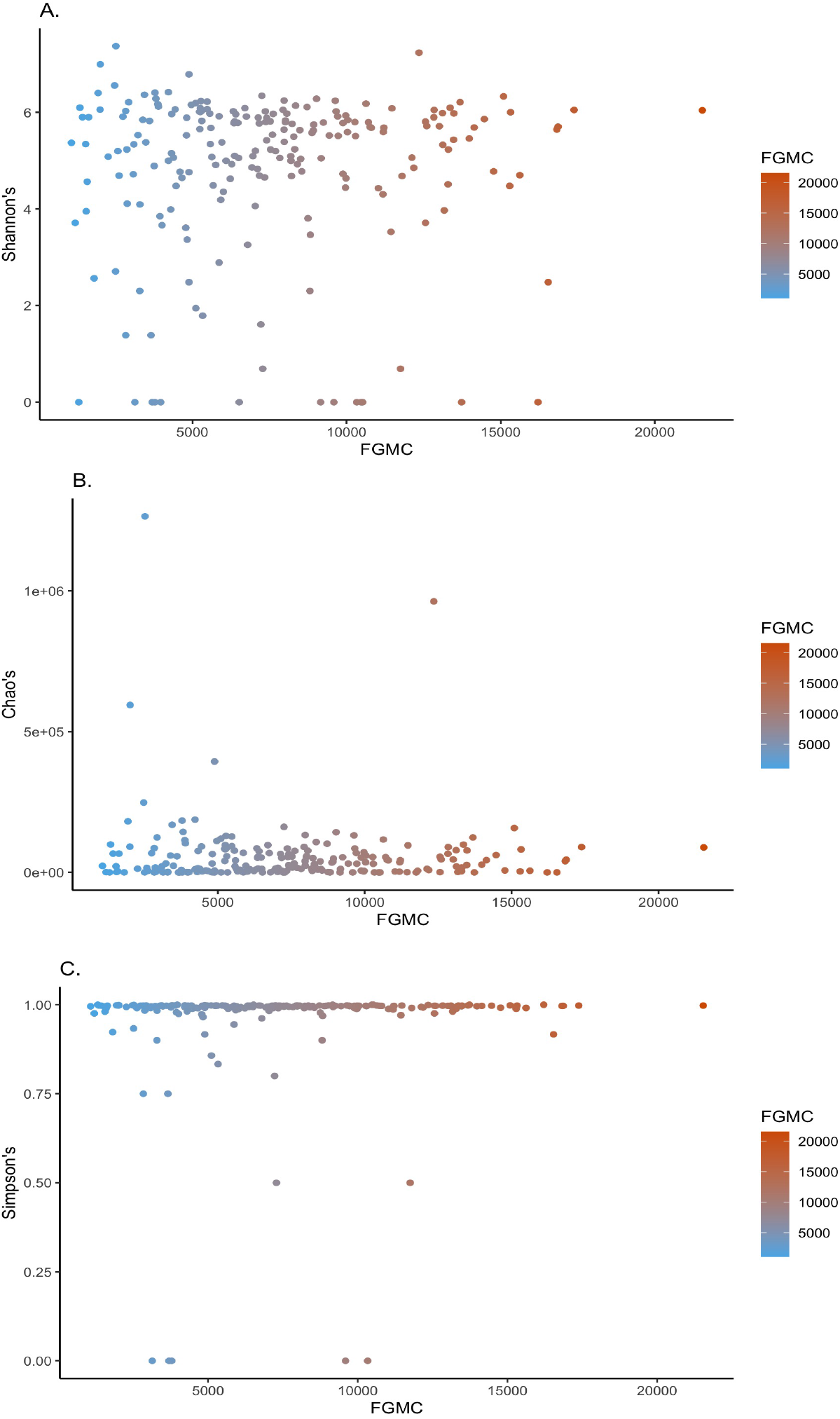
Scatterplots of FGMC against A. Shannon’s diversity and B. Chao’s diversity index were not significant

**Figure 2.**
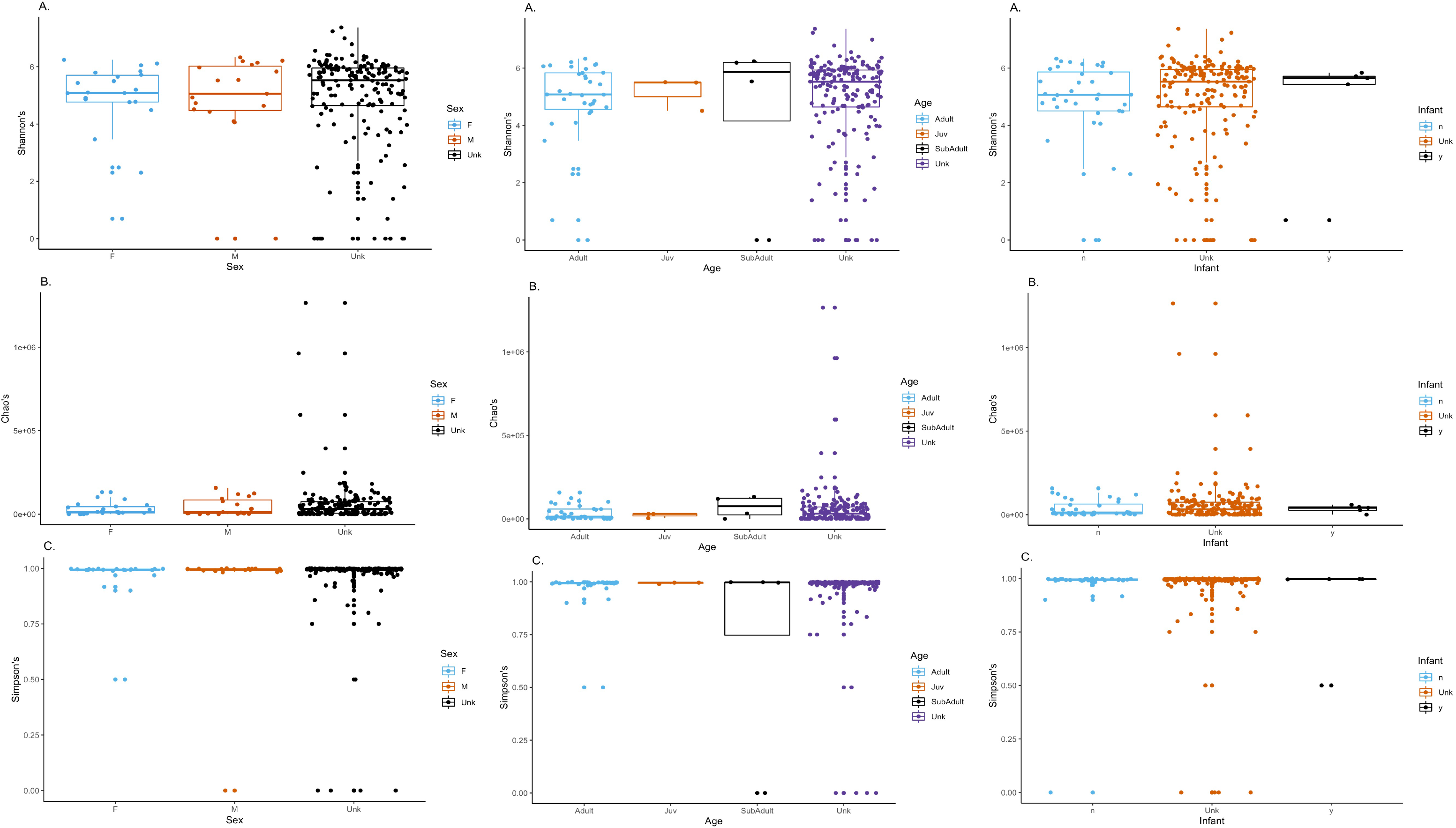
The predictor variables, sex (column 1), age (column 2), and infant (column 3) were all not significant for A. Shannon’s and B. Chao’s.

Bonobo’s FGMC did not change due to sex, age, and whether or not a female had an infant. We found no significant interaction between FGMC and sex (Shannon: p-value = 0.131; Chao’s: p-value = 0.510), age (Shannon: p-value = 0.143; Chao’s: p-value = 0.459), and whether or not a female had an infant (Shannon: p-value = 0.131; Chao’s: p-value = 0.487) for both alpha diversity metrics when run in two-way ANOVA with an interaction effect. FGMC values had mean 7529 ng/pl ± 266.55 ng/pl and did not significantly differ by sex (p-value = 0.387), age (p-value = 0.17), and whether or not a female had an infant (p-value = 0.144).

### Beta diversity

Bonobo FGMC significantly explained a small amount of variation in beta diversity for two of the three beta diversity metrics, Jensen-Shannon’s Distance and weighted UniFrac. Beta diversity did not significantly relate to FGMC for unweighted UniFrac. The PERMANOVA results for FGMC showed that it explained 0.08% of the variation in beta diversity for the Jensen-Shannon Distance (Table 2). The PERMAOVA for the weighted UniFrac dissimilarity matrix found FGMC explained 1.27% (Figure 3; Table 2). The PERMANOVA results for FGMC showed that the unweighted UniFrac dissimilarity matrix did not significantly explain variation in beta diversity, but it was suggestive of being significant at a p-value = 0.06 (Table 2).

**Figure 3.**
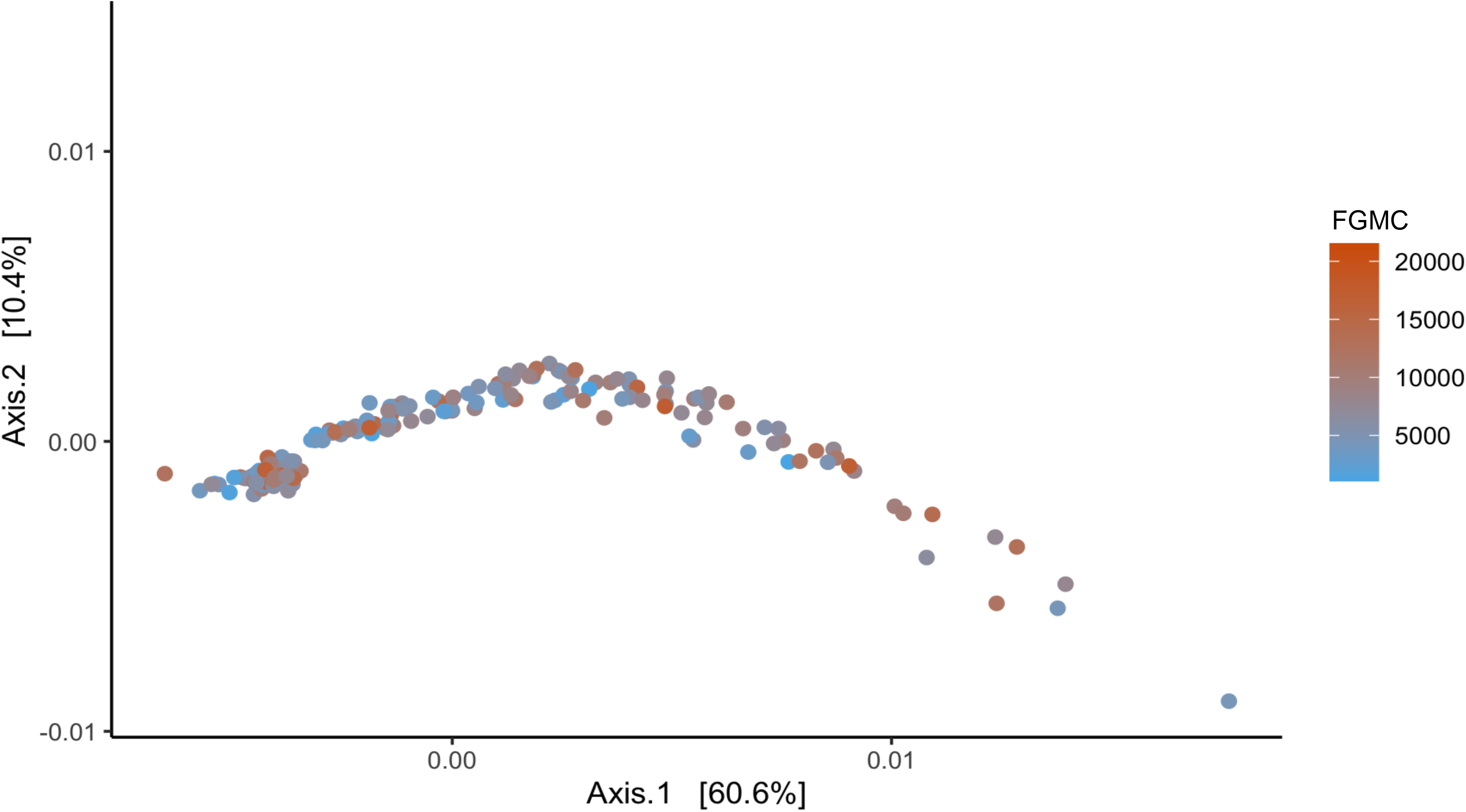
PCoA plot of weighted UniFrac dissimilarity matrix. The PERMANOVA results suggest that FGMC explained 1.275% of the variation in beta diversity. While, weighted UniFrac explained the most of the variation in beta diversity, this is very small amount of the variation in beta diversity.

**Table 2.**
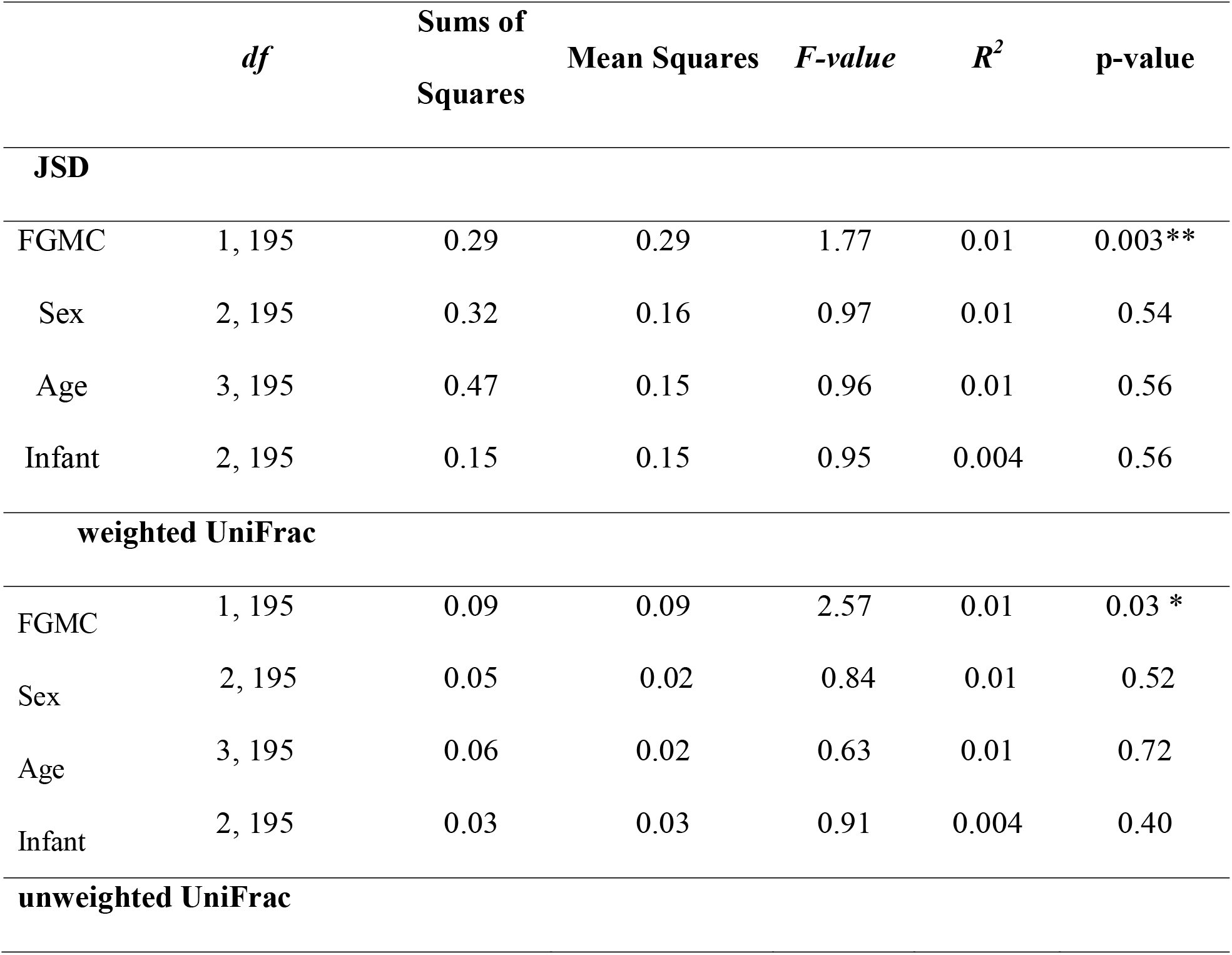
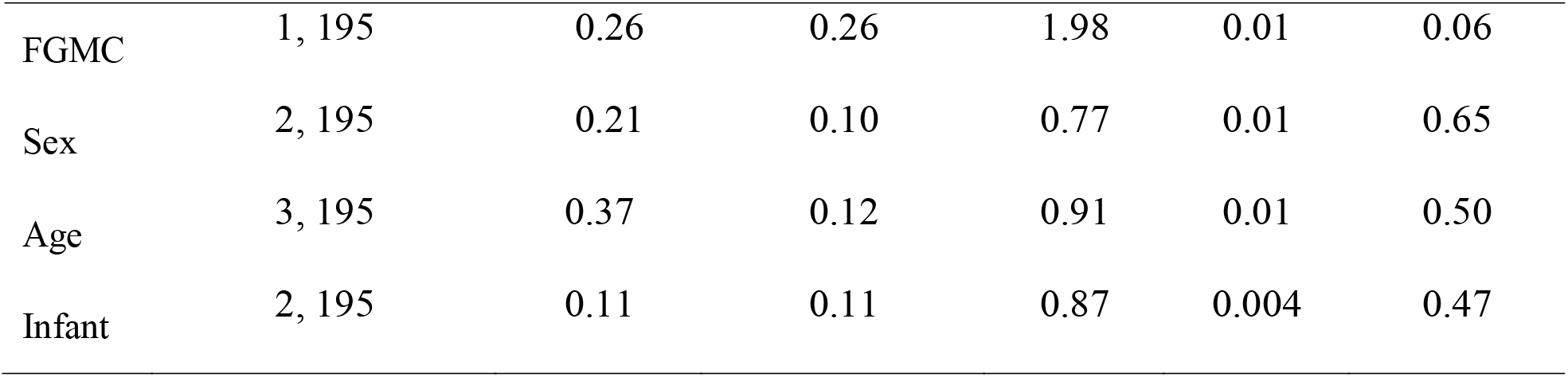
PERMANOVA results for Jensen-Shannon Distance (JSD), weighted UniFrac, and unweighted UniFrac against FGMC, sex, age, and whether or not a female had an infant.

Mantel test showed that there was a significant agreement between the FGMC Euclidian distance matrix and the Jensen-Shannon Distance matrix showing that large difference in FGMC value were associated with high values of dissimilarity in Jensen-Shannon Distance (Observed value: 0.09, p-value = 0.02) (Figure 4). The Mantel test for weighted UniFrac (p-value = 0.28) and unweighted UniFrac (p-value =0.17).

**Figure 4.**
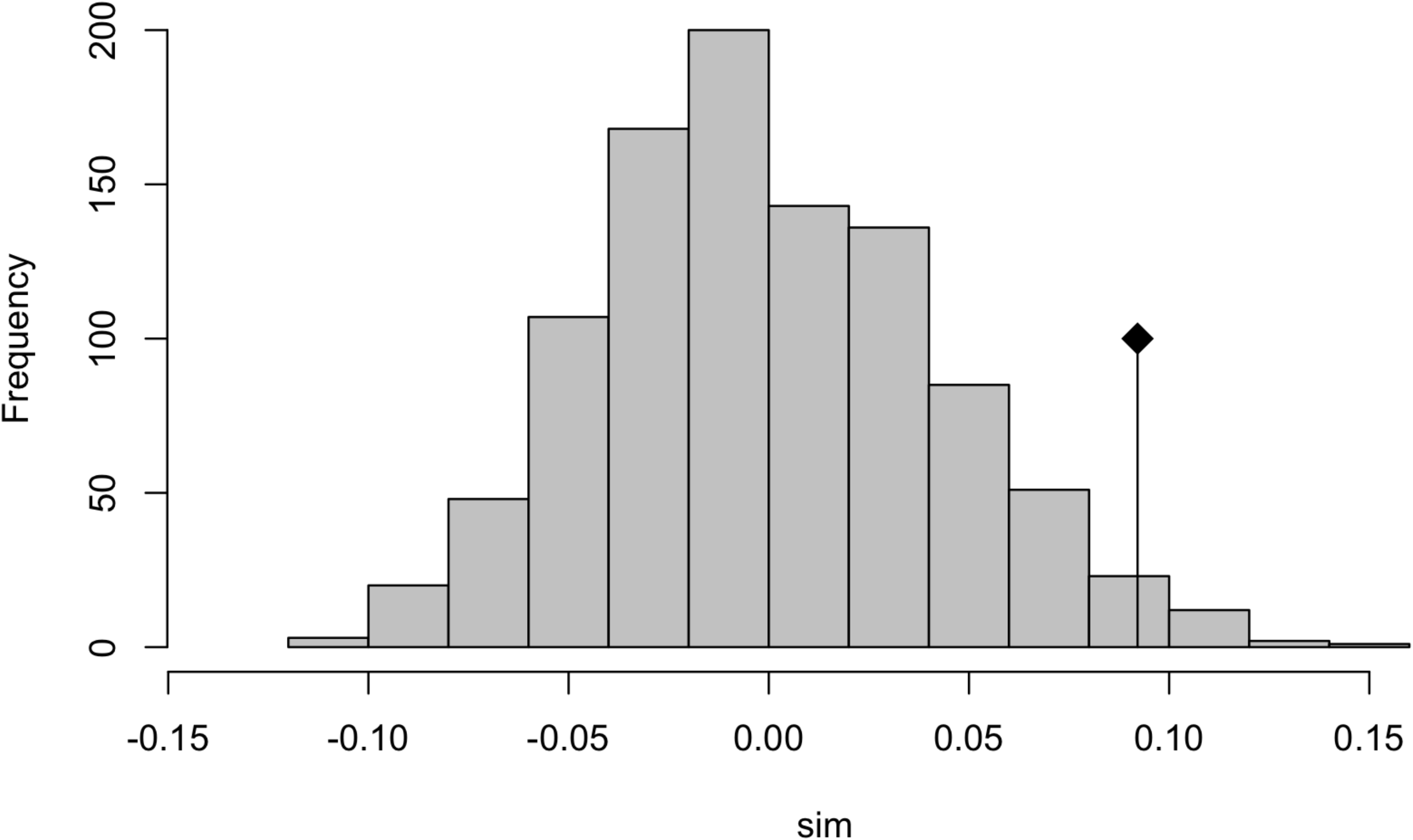
Frequency distribution from the Mantel test showing the results of the 999 randomizations for Jensen Shannon Distance matrix and the FGMC Euclidian distance matrix. The observed value is marked with the line and black diamond.

### Analysis of community variance (ANCOM) results

Bonobos have 17 taxa whose abundance was explained by FGMC after Bonferroni correction out of 302 taxa tested (Table 3; Supplementary Figure 1). We found 15 of the significant ASVs had a positive linear relation with FGMC while two significant ASVs *RFN20 and Butyrivibrio*, had a negative linear relation with FGMC.

**Table 3.**
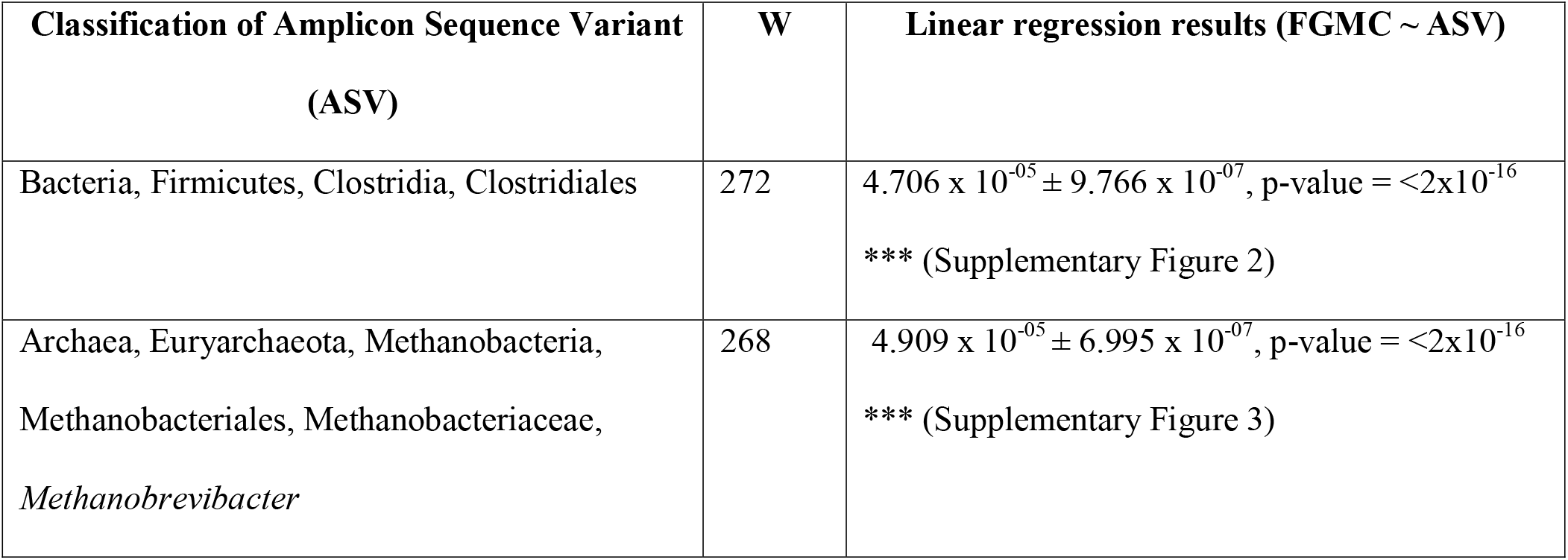
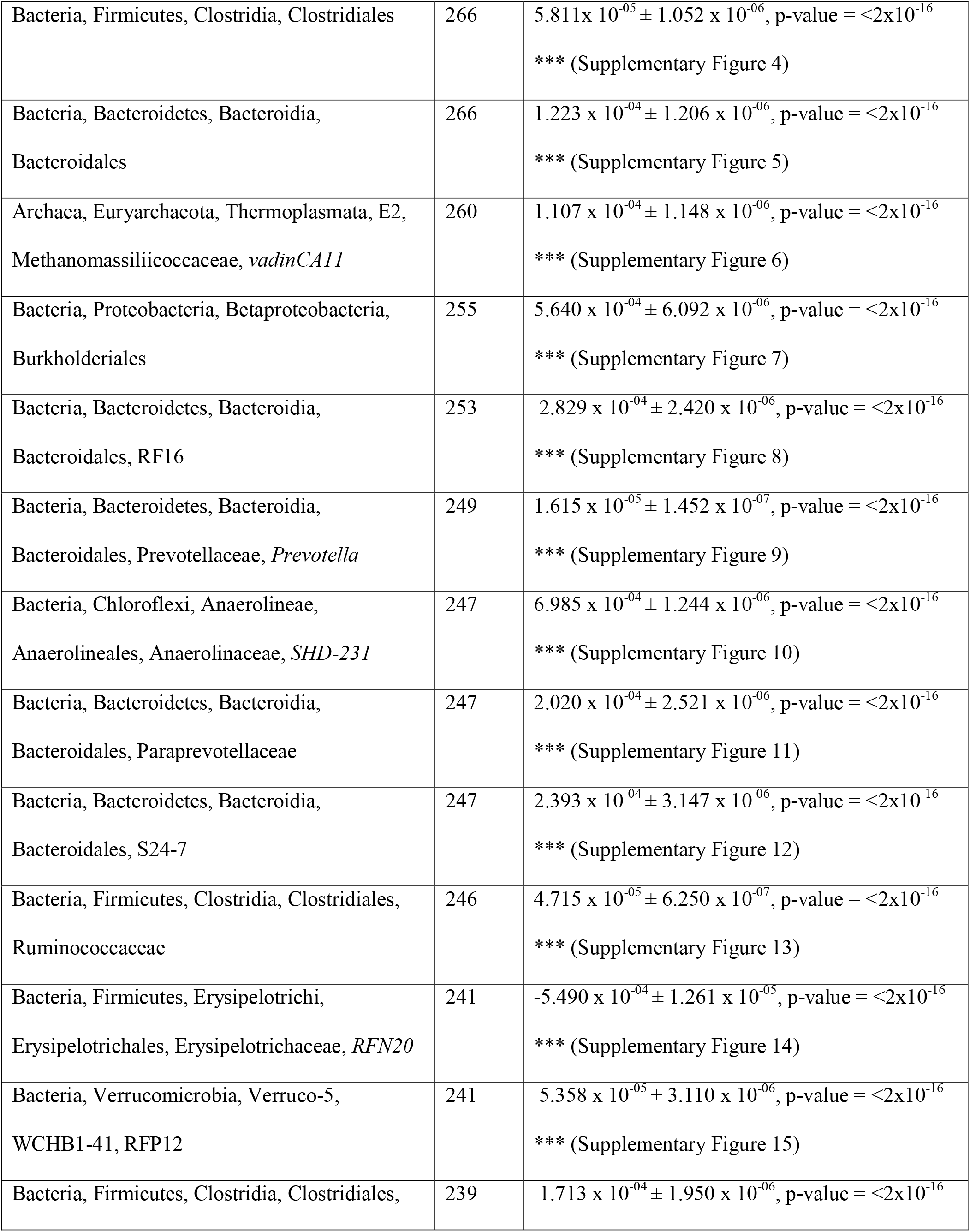
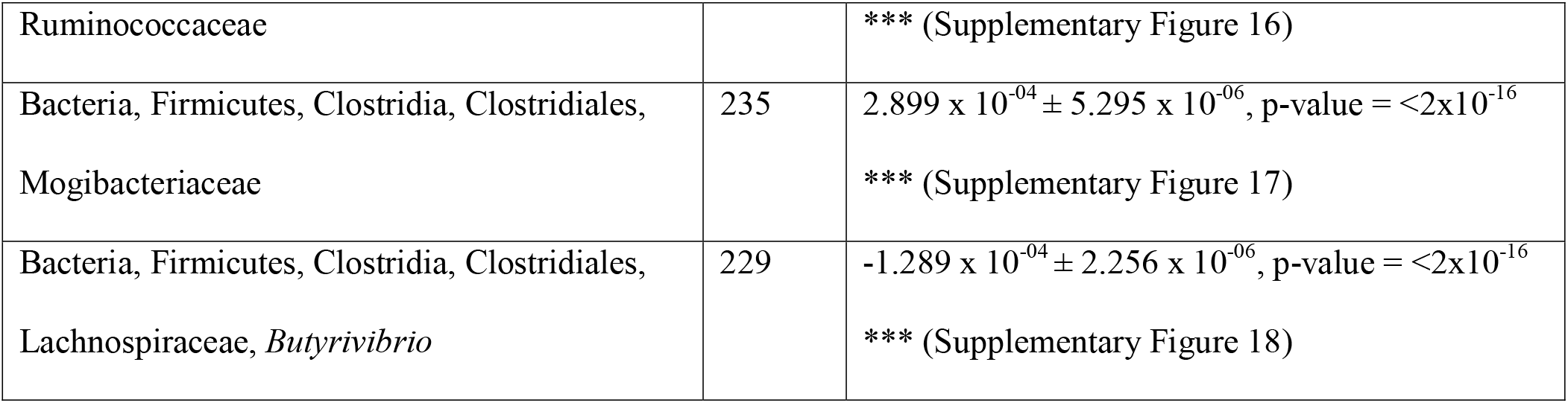
Taxa from ANCOM results were significantly related to FGMC after Bonferroni correction. W is equal to the number of times the log-ratio of a particular taxon compared to every other taxon being tested was detected to be significantly different across groups. For all the listed taxa, we accepted the alterative hypothesis that the FGMC does significantly explain the abundance of the listed taxa. The taxa are listed by: Kingdom, Phylum, Class, Order, Family, Genus.

## Discussion

We aimed to test the relationship between FGMC and the gut microbiota for bonobos and compare the results to those reported for western lowland gorillas, humans, and other mammals [10]. We predicted that the bonobo gut microbiota would exhibit patterns similar to what was found in western lowland gorillas, where alpha diversity (or within individual diversity) and beta diversity (or between individual diversity) were not significantly explained by FGMC due to the similar social environment, ecology, and phylogenetic relationship between western lowland gorillas and bonobos. The abundance model results for the western lowland gorillas found three taxa significantly correlated with FGMC [10].

Consistent with Vlčková et al. (2018), our alpha diversity measures were not significant. This suggests that number of bacterial taxa within the gut microbial community stays constant. Additionally, it suggests there is stability in the number of taxa in the guts of great apes even when the gut microbiome is disrupted by stress. This is constant with several other finding outside of great apes, where despite potentially stressful habitat fragmentation, red colobus (*Procolobus rufomitratus*), black and white colobus (*Colobus guerza*), and red-tailed guenon (*Cercopithecus ascanius*) gut microbiomes remained stable [90]. This stability in alpha diversity may be a wild primate feature as other mammals like elephants [31], pangolins [30], and squirrels [91] have been found to exhibit changes in alpha diversity linked to FGMC. This may point to the ability of primates to buffer stressful events due to their behavioral flexibility and social relationships [92], [93].

Our beta diversity results found that FGMC significantly explains a small amount of between individual variation in the bonobo gut microbiota. The amount of variation explained in our PERMANOVA by FGMC was very small and could be due noise in the data, but because we also detected a difference with the mantel tests for the JSD dissimilarity metric this result is likely an actual pattern. The small amount of variation explained implies that there is only a small number of taxa whose abundance is affected by FGMC. In other primates, beta diversity has been found to significantly change with degraded habitats in howler monkeys (*Alouatta pigra*), potentially due to the stress of inhabiting and eating much lower quality food items [94]. In contrast, beta diversity for red colobus (*Procolobus rufomitratus*), black and white colobus (*Colobus guerza*), and red-tailed guenon in degraded habitats remained stable [90]. In primates, there doesn’t seem to be a clear pattern of how stress and beta diversity relate to each other and may depend on the context that a wild primate is living in or may depend on the specific taxa found in the gut of a wild-living primate. In mammals, elephants [31]and pangolins [30], beta diversity significantly changes with a stress. Beta diversity changes in response to stress appear to be context and species specific. The fact that beta diversity was significant for bonobos and not gorillas implies that bonobos gut microbiomes may be more susceptible to stress or that bonobo gut microbiomes are home to bacteria whose abundance is more susceptible to fluctuations in stress. Our abundance model results support the idea that bonobos house more taxa whose abundance changes with fluctuations in stress.

There were fourteen more taxa in the bonobo gut microbiome whose abundance was significantly related to FGMC, for both our ANCOM and linear model results. Vlčková et al. (2018) found only three microbial taxa were shown to be significant with FGMC in western lowland gorillas (family Anaerolineaceae; genus *Clostridium* cluster XIVb; genus *Oscillibacter)* [10]. We found members of the family Anaerolineaceae in bonobo samples, and interestingly one taxon (genus SHD-231) was significant in our differential abundance model and linear model results, similar to gorillas [10]. We did find one genus of *Clostridium* in the 201 bonobos that we sampled; however, this genus of *Clostridium* was not a part of cluster XIVb, found in western lowland gorillas to be significant [10]. We found no genus *Oscillibacter,* a genus found to be significant gorillas, in our bonobo samples, nor did we detect any of the higher family level Oscillospiraceae. Therefore, bonobos and western lowland gorillas appear to be similar in that members of the family Anaerolineaceae may be differentially affected during periods of high stress but differ in the sixteen other taxa that are differentially abundant based on FGMC in bonobos.

Other taxa that we found specific to the bonobo abundance results are two different unknown bacteria in the order Clostridiales that are thought to be linked to early life stress in mice [95]. Additionally, two members of the order Bacteroidales, including family S24-7, which has been associated with changes in circadian rhythm disruption in murine models [96]. We also found Ruminococcaceae and Mogibacteriaceae, which have been found in the human gut microbiota [97], [98]. Other notable genus level associations included *Prevotella* sp., associated with chronic inflammatory conditions [99]. These different patterns between the bonobo and western lowland gorillas suggest that there may be species-specific or temporal-specific effects of FGMC on the abundance of specific taxa in primate gut microbiota, and several of these taxa have been linked to early life stress, circadian rhythm stress, and chronic inflammation [27], [95], [96]. However, it does appear that the family Anaerolinaceae may be particularly affected by stress in great apes.

In other mammals, like squirrels, pangolins, elephants, and rhinos, stress was associated with changes, in specific taxa [30]–[32], [91]. Additionally, specific taxa changed in abundance due to stress or stressful events [28], [30], [32]. The specific taxa are differentially expressed in elephants (Planococcaceae, Clostridiaceae, Spirochaetaceae, and Bacteroidia) and rhinos (Aerococcaceae, Atopostipes, Carnobacteriaceae, and Solobacterium) were not found to be related explicitly to FGMC in bonobos [31]. However, there may be high order similarities between bonobos and other mammals in the specific taxa that are differential abundant due to stress. Clostridiaceae is a family belonging to the class Clostridia that was found to be differentially expressed in elephants [31]. While we did not find this specific family differential expressed in bonobos, we found six Clostridia members to be differentially expressed in bonobos. This result may indicate that stress may affect specific taxa in the gut microbiota across mammalian lineages, including humans. In humans, the proposed mechanism for a host and its associated microbes to communicate is through hormones [15], [81]. We did not see an overall decrease in diversity as has been seen in humans, but we did see certain taxa whose abundance seems to be differentially affected by stress [9]. Indicating that there may be similarities between humans and one of their closest evolutionary relatives, bonobos.

There are several differences in methods that could be influencing stress and the gut microbiota between bonobos and gorillas. One of the significant differences between our study and the gorilla study is the sample size. Our larger sample size may be why we picked a small effect of beta diversity and stress, but more studies across wild living primates examining stress and the gut microbiota will help to elucidate how sample size plays into picking up relationships between proximate measures of stress and the gut microbiota. Other differences include, potentially different FGMC measures and differences in the method of sample preservation for FGMC analysis [10], [100]. There are also species-specific patterns in the production, metabolism, and excretion of FGMCs [78]. Therefore, comparing FGMC values between bonobos and gorillas must be done with extreme care. However, since we are not directly comparing our FGMC values to those obtained for the western lowland gorillas. Instead, we are comparing the broad results from comparing those FGMCs to the gut microbiota. Additionally, the gut microbiota may be responding to FGMCs in non-linear ways, and there may be more nuanced changes to consider when comparing stress and the NHP gut microbiota.

Other limitations include the metabolism of FGMCs can also be dependent on sex and time of day [79], [101]. At the same time, we attempted to control for this variation by only selecting samples collected around the same time of day and including sex and age as factors in our analyses. This method of controlling for time of day is like the western lowland gorilla paper, where morning fecal samples were analyzed for the unhabituated groups [10]. In addition to variation in FGMCs, there are several other factors that influence the composition and diversity of gut microbiota among NHP.

There are other variables that we did not examine in this paper that have been thought to influence the composition and diversity of the gut microbiota in NHP. Disease status could be influencing the gut microbiota [14]. Rank and other social factors like rates of affiliation and aggression both within and between communities could also be affecting the gut microbiota [74], [102], [103]. Diet and seasonality can also be significant factors in changing how nutritionally stressed an individual is and can directly affect the gut microbiota composition [104]–[106]. We aim to examine these factors in future analyses.

Future directions for this work include adding metagenomic sequencing and metabolomic data to our dataset to incorporate more functional results. Additionally, we aim to incorporate diet, food availability, and social variables in future analyses of the bonobo gut microbiota. Compared to humans, in bonobos, beta diversity and some taxa change in abundance instead of broadly changing gut microbial diversity in response to stress, but we did find similarities in bonobos and gorillas in the family Anaerolinaceae. Thus, future studies should examine how Anaerolinaceae changes in response to stress in other great apes like humans, chimpanzees, and orangutans. Incorporating FGMC and gut microbiota data can provide a more robust understanding of how stress impacts the gut microbiota of primates.

## Acknowledgments

We want to thank the following people without whom this work would not be possible: Jef Dupain, Hugues Akpona, and the African Wildlife Foundation, Dipon Bomposo, Beken Bompoma, Teddy Bofaso, Christian Djambo, Mathieu Esaola, Bellevie Iyambe, Augustin Lofili, Isaac Lokoli, Gedeon Lokofo, Thomas Lokuli, Joel Bontambe, Abdulay Bokela, Papa Siri, Paco Bertolani, Ian Takaoka, and many others who generously provided both logistical support and friendship in the field. Additionally, we want to thank Diana Christie and the Molecular Anthropology laboratory group members for their helpful discussions through all stages of this research.

## Statement of Ethics

This research was completed with the approval of the University of Oregon’s Institutional Animal Care and Use Committee (AUP-17-10), abided by the American Society of Primatologist’s Principles for the Ethical Treatment of Human Primates, and was approved by the L’Institut Congolais pour la Conservation de la Nature (ICCN) (permit # 3058) the department withing the Democratic Republic of the Congo that oversees the Lomako-Yokokala Faunal Reserve where the Iyema field site is located. Permissions were obtained through the African Wildlife Foundation Mission Order (No. 033/2017).

## Author Statements

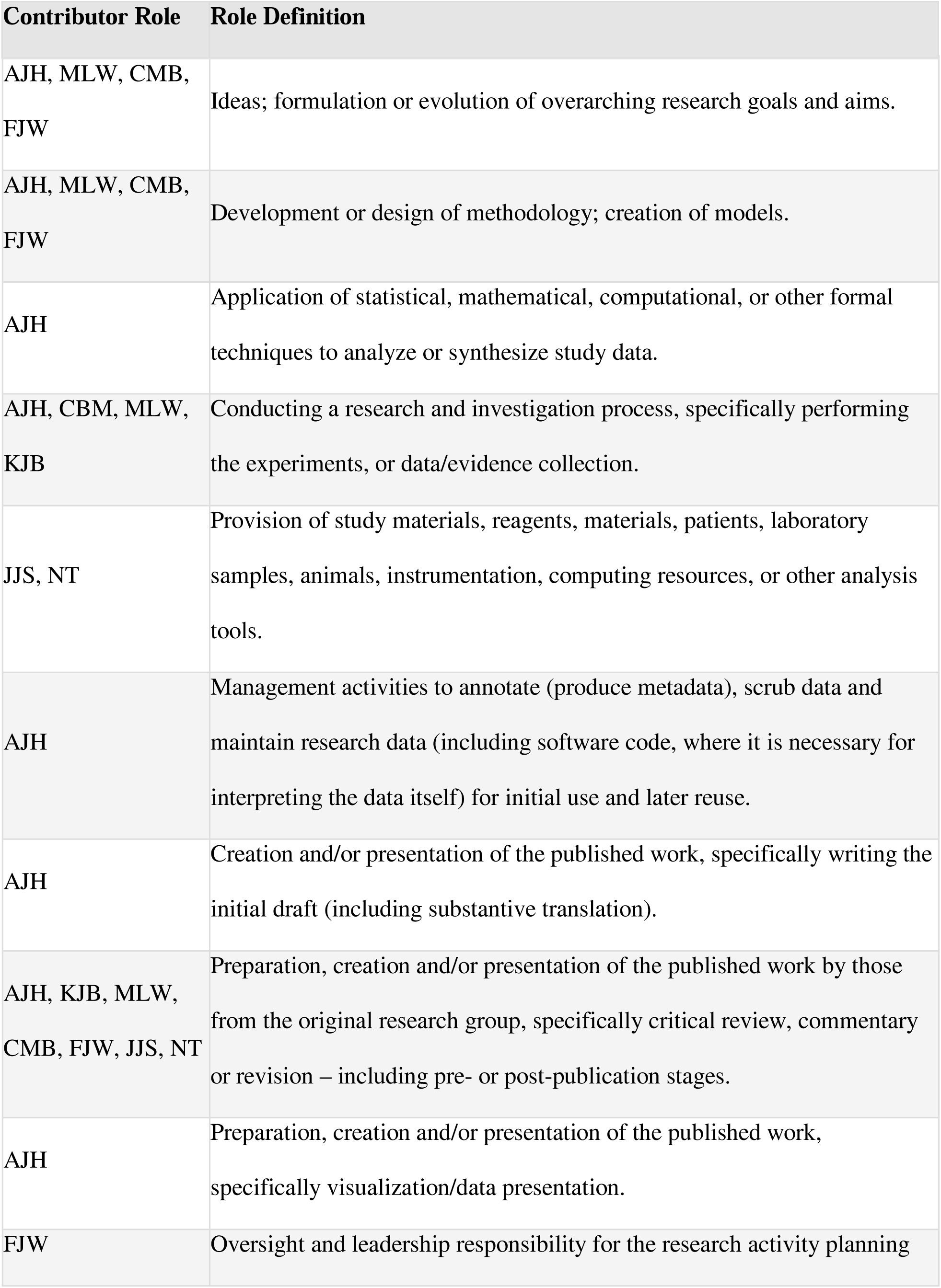
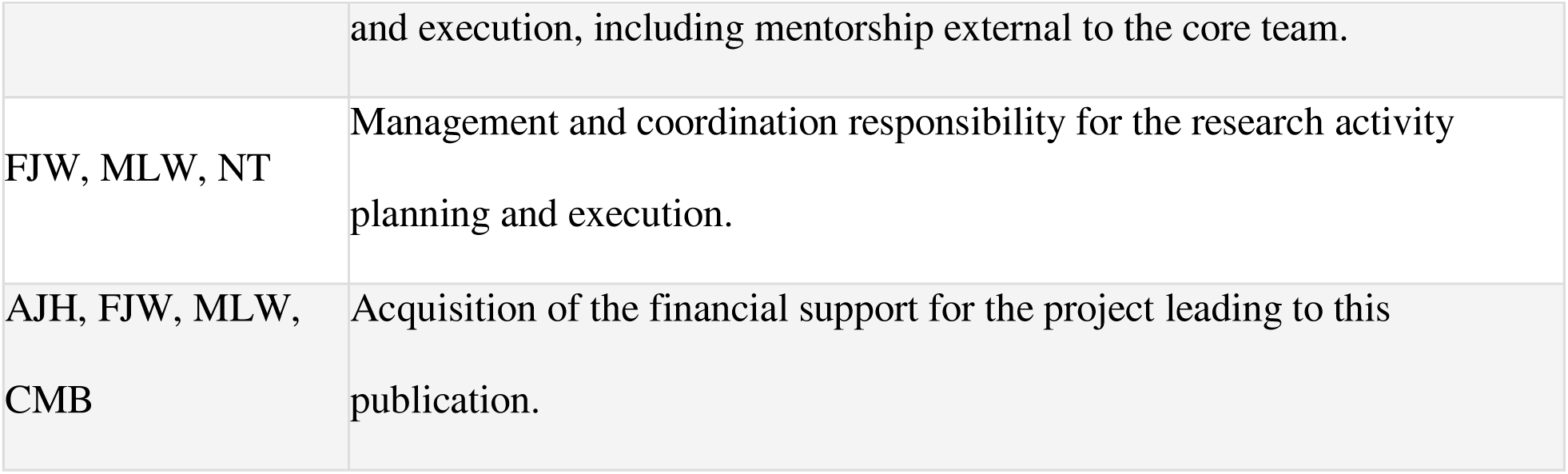

---

***Supplementary Figure 1.** Volcano plot of W and F statistic for each ASV. 17 taxa (circled in blue) are significantly related to FGMC level*.

***Supplementary Figure 2.*** General linear model regression on the ASV Bacteria, Firmicutes, Clostridia, Clostridiales and FGMC.

***Supplementary Figure 3.*** General linear model regression on the ASV Archaea, Euryarchaeota, Methanobacteria, Methanobacteriales, Methanobacteriaceae, *Methanobrevibacter* and FGMC.

***Supplementary Figure 4.*** General linear model regression on the ASV Bacteria, Firmicutes, Clostridia, Clostridiales and FGMC.

***Supplementary Figure 5.*** General linear model regression on the ASV Bacteria, Bacteroidetes, Bacteroidia, Bacteroidales and FGMC.

***Supplementary Figure 6.*** General linear model regression on the ASV Archaea, Euryarchaeota, Thermoplasmata, E2, Methanomassiliicoccaceae, *vadinCA11* and FGMC.

***Supplementary Figure 7.*** General linear model regression on the ASV Bacteria, Proteobacteria, Betaproteobacteria, Burkholderiales and FGMC.

***Supplementary Figure 8.*** General linear model regression on the ASV Bacteria, Bacteroidetes, Bacteroidia, Bacteroidales, RF16 and FGMC.

***Supplementary Figure 9.*** General linear model regression on the ASV Bacteria, Bacteroidetes, Bacteroidia, Bacteroidales, Prevotellaceae, *Prevotella* and FGMC.

***Supplementary Figure 10.*** General linear model regression on the ASV Bacteria, Chloroflexi, Anaerolineae, Anaerolineales, Anaerolinaceae, *SHD-231* and FGMC.

***Supplementary Figure 11.*** General linear model regression on the ASV Bacteria, Bacteroidetes, Bacteroidia, Bacteroidales, Paraprevotellaceae and FGMC.

***Supplementary Figure 12.*** General linear model regression on the ASV Bacteria, Bacteroidetes, Bacteroidia, Bacteroidales, S24-7 and FGMC.

***Supplementary Figure 13.*** General linear model regression on the ASV Bacteria, Firmicutes, Clostridia, Clostridiales, Ruminococcaceae and FGMC.

***Supplementary Figure 14.*** General linear model regression on the ASV Bacteria, Firmicutes, Erysipelotrichi, Erysipelotrichales, Erysipelotrichaceae, *RFN20* and FGMC.

***Supplementary Figure 15.*** General linear model regression on the ASV Bacteria, Verrucomicrobia, Verruco-5, WCHB1-41, RFP12 and FGMC.

***Supplementary Figure 16.*** General linear model regression on the ASV Bacteria, Firmicutes, Clostridia, Clostridiales, Ruminococcaceae and FGMC.

***Supplementary Figure 17.*** General linear model regression on the ASV Bacteria, Firmicutes, Clostridia, Clostridiales, Mogibacteriaceae and FGMC.

***Supplementary Figure 18.*** General linear model regression on the ASV Bacteria, Firmicutes, Clostridia, Clostridiales, Lachnospiraceae, *Butyrivibrio* and FGMC.

***Supplementary Figure 19.*** General linear model regression on the first principal coordinate and FGMC.

