## Supplementary figures and images for "A comparison of fecal glucocorticoid metabolite concentration and gut microbiota diversity in bonobos (*Pan paniscus*)"

### Supplementary figure 1

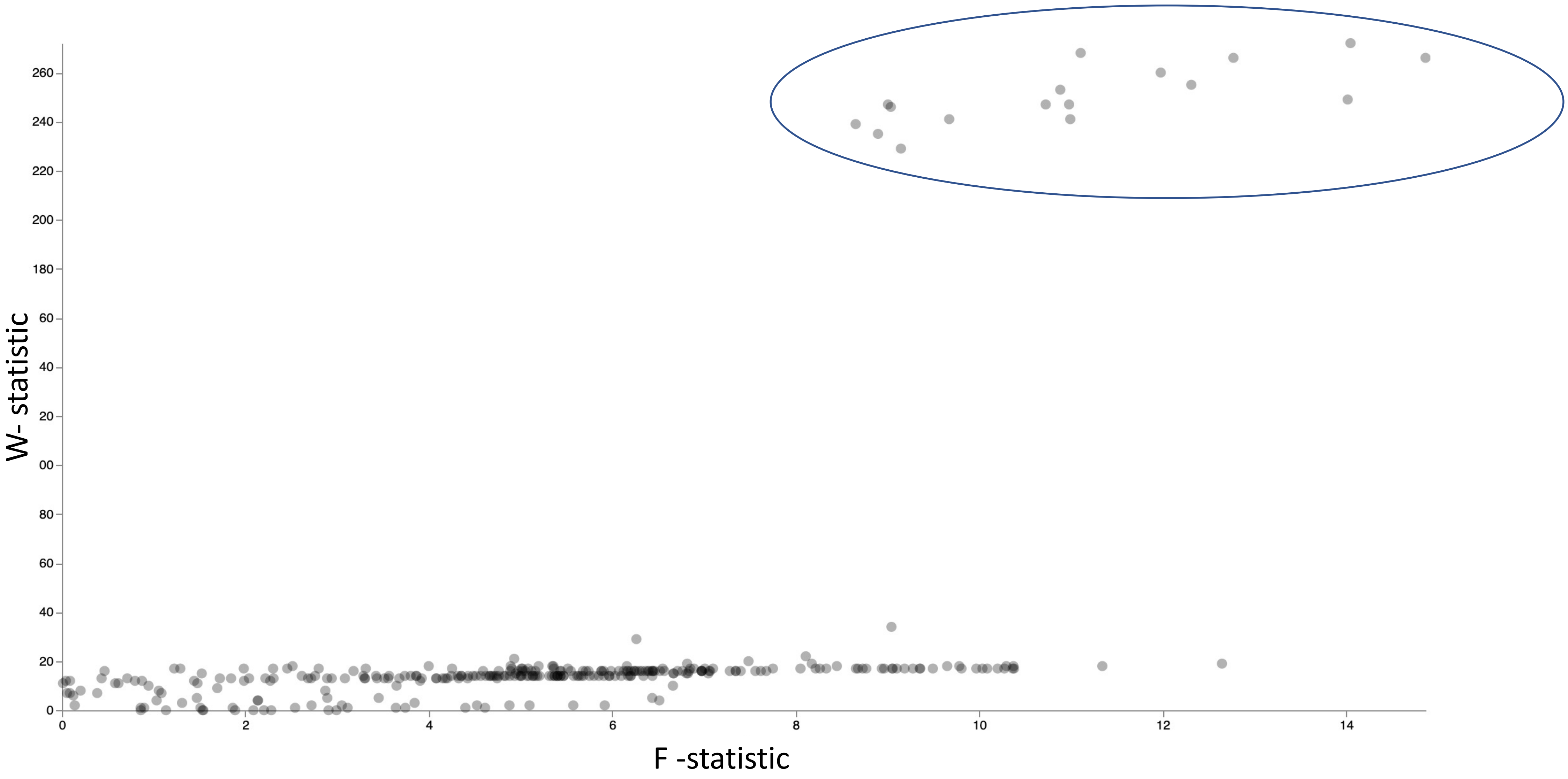

### Supplementary figure 2

**Supplementary Figure 2:**

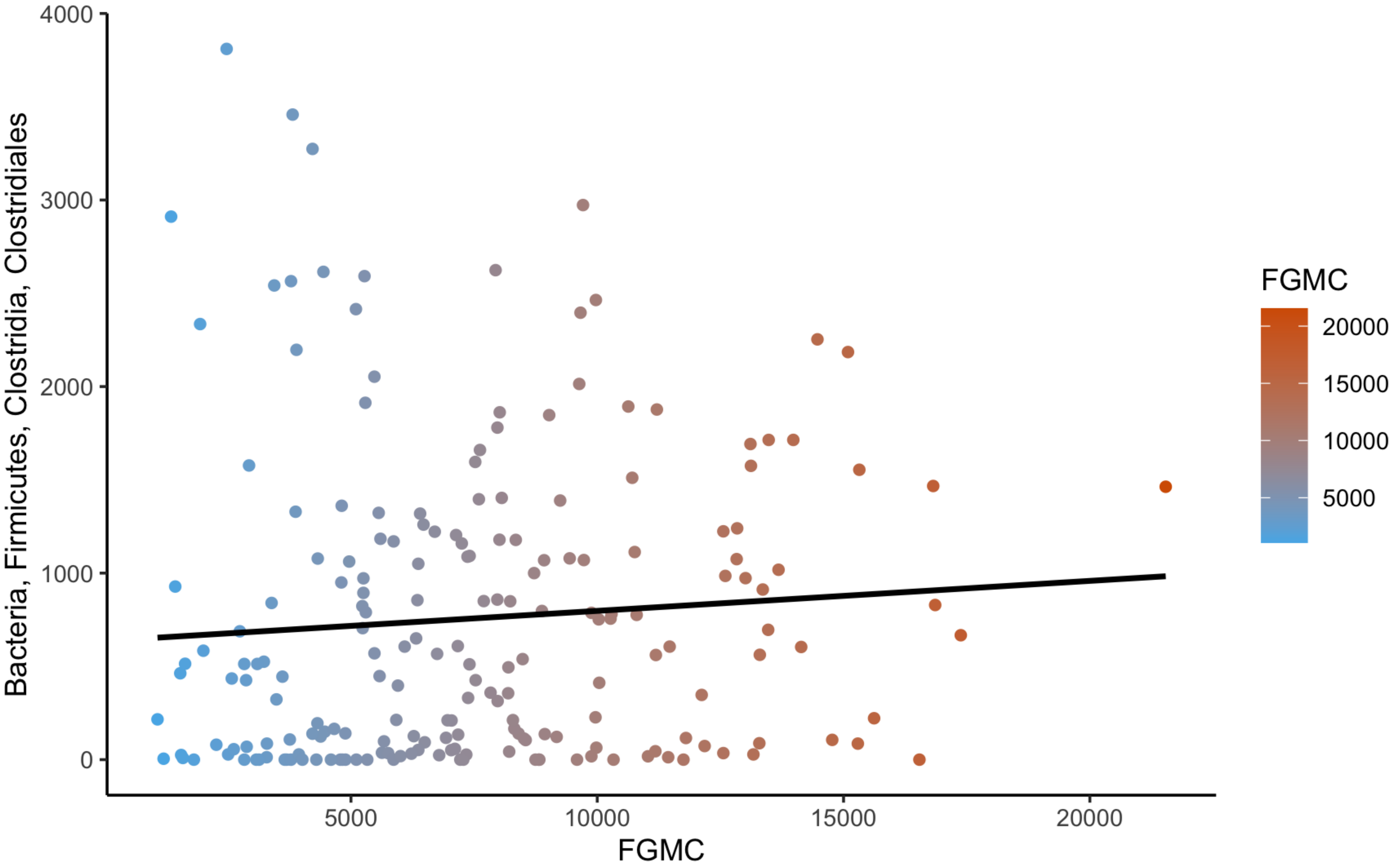

### Supplementary figure 3

**Supplementary Figure 3:**

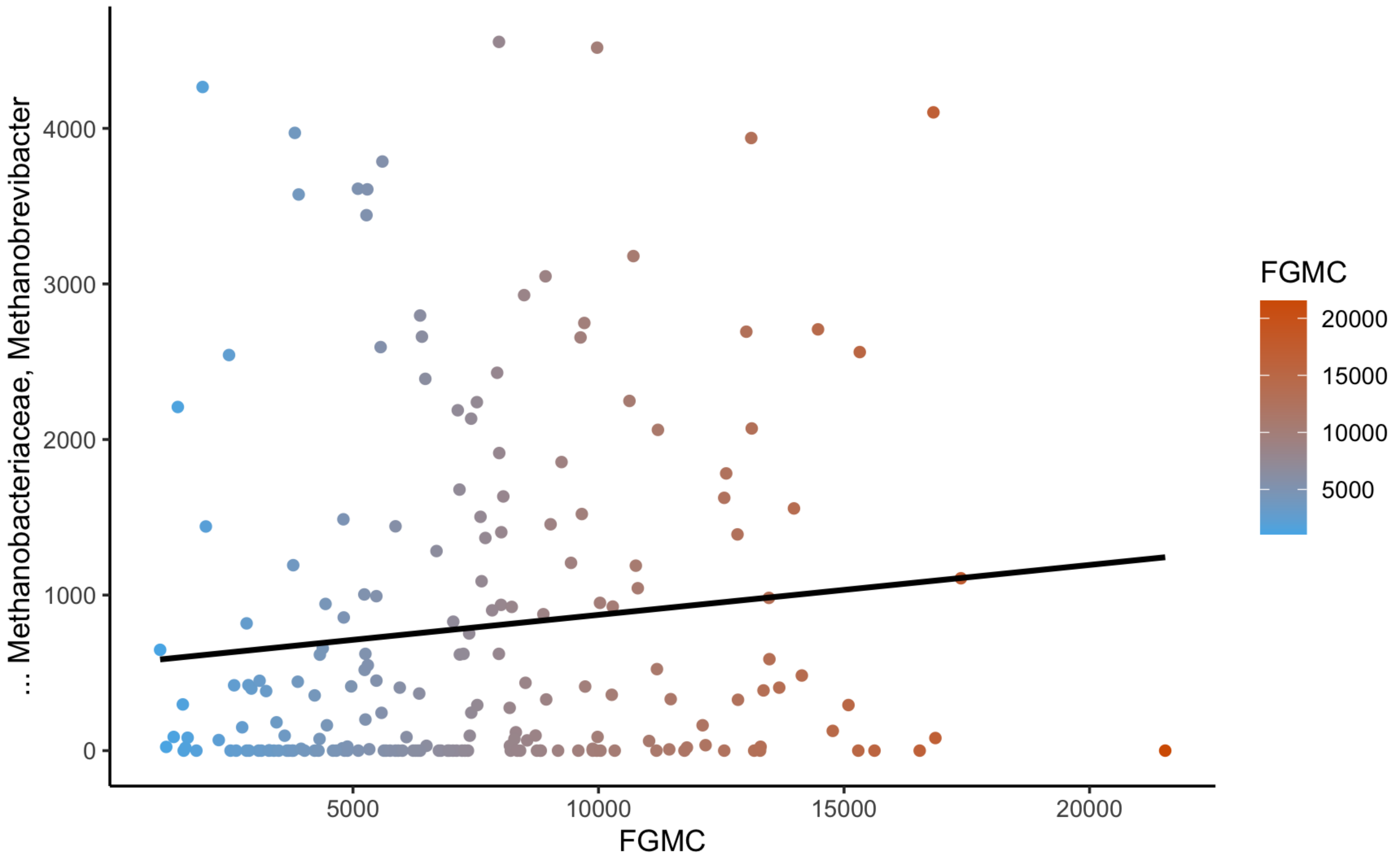

### Supplementary figure 4

**Supplementary Figure 4:**

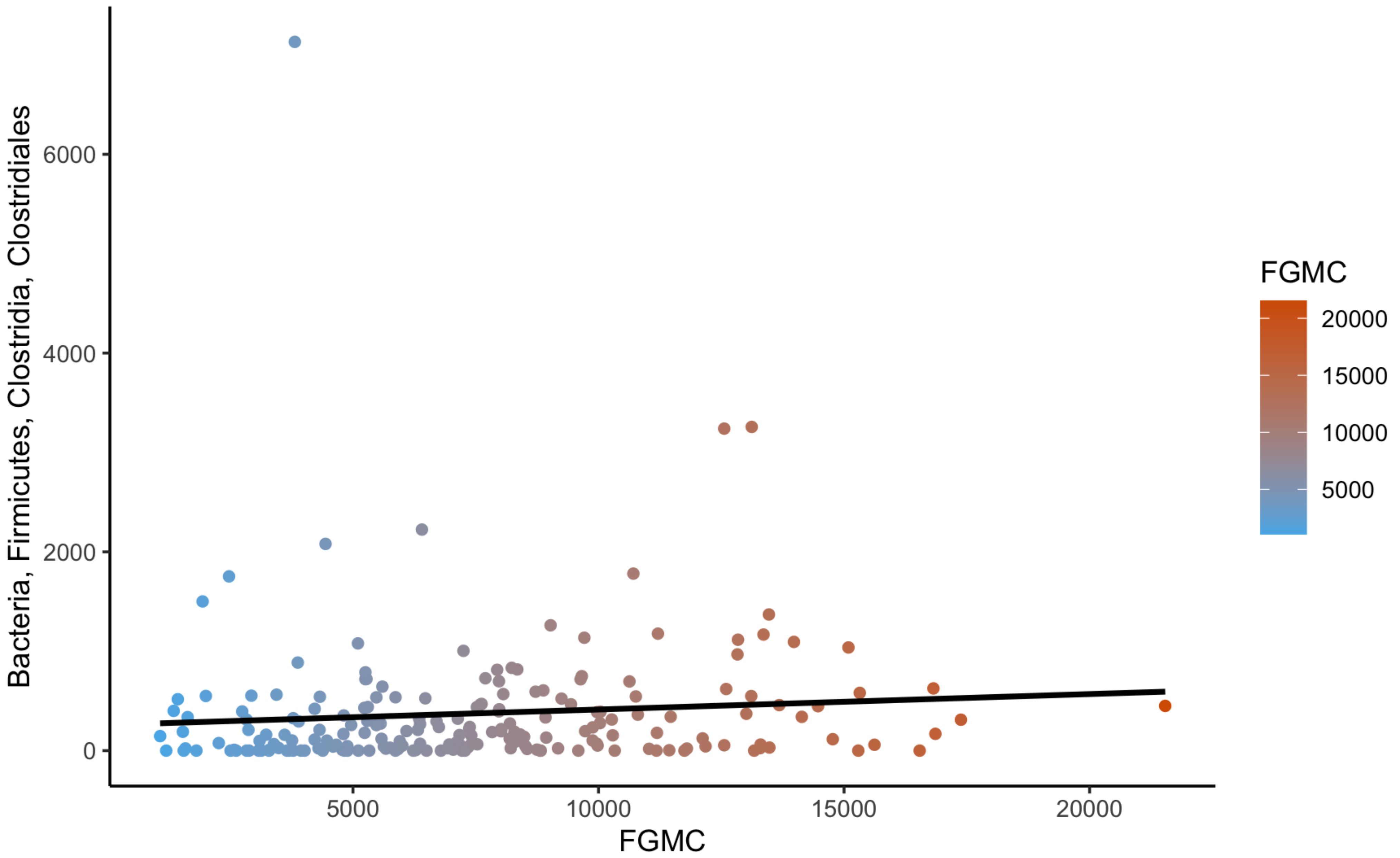

### Supplementary figure 5

**Supplementary Figure 5:**

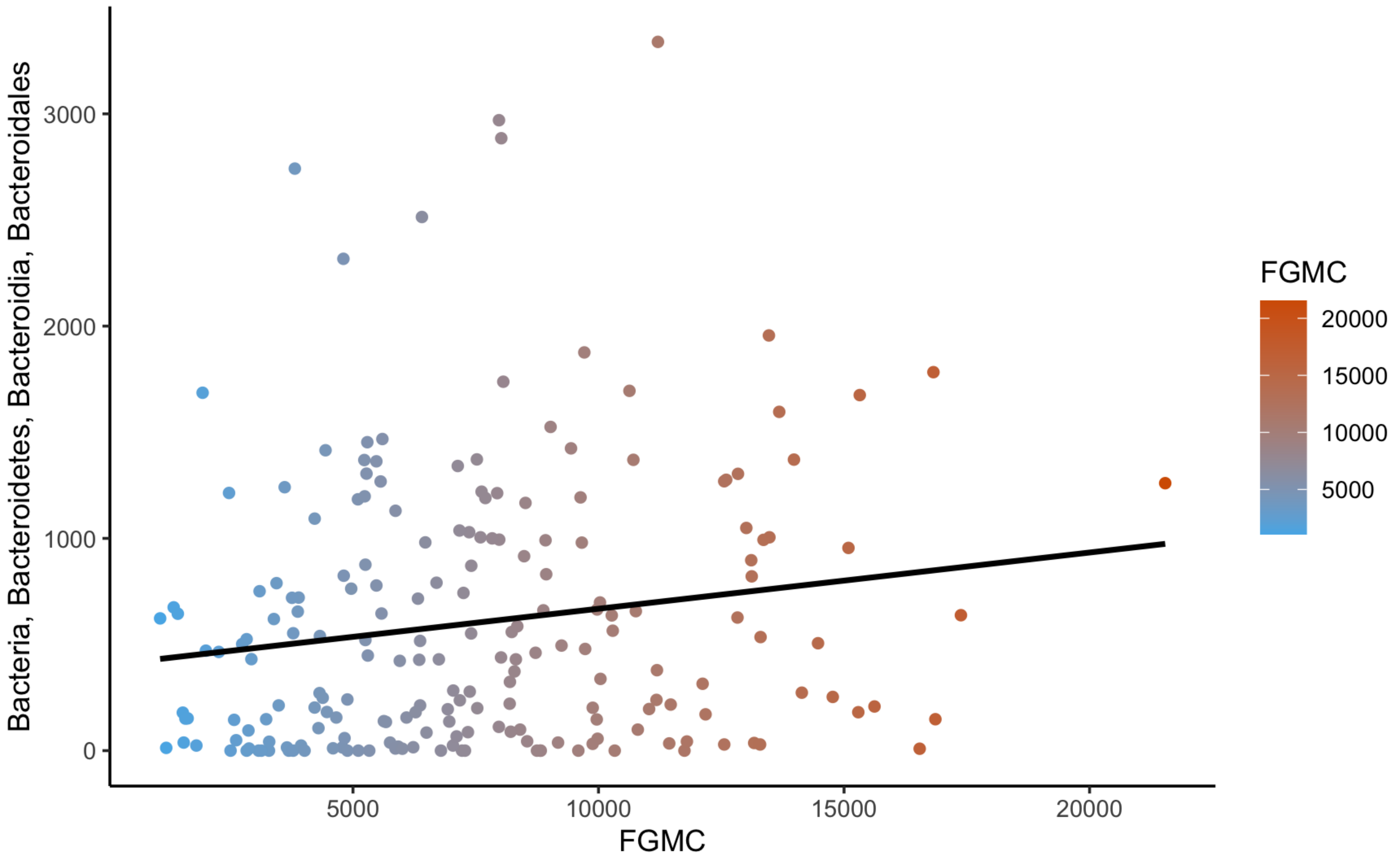

### Supplementary figure 6

**Supplementary Figure 6:**

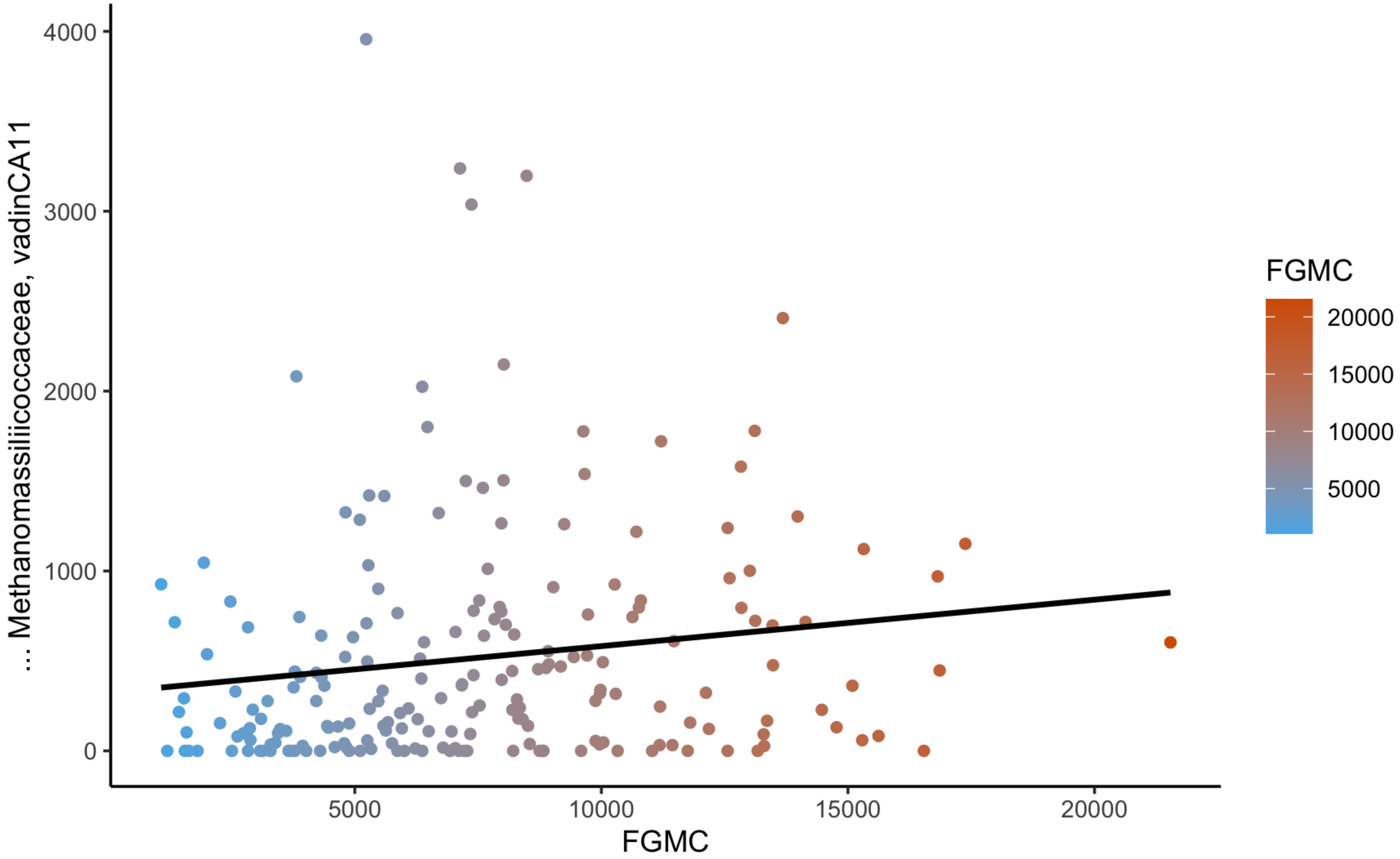

### Supplementary figure 7

**Supplementary Figure 7:**

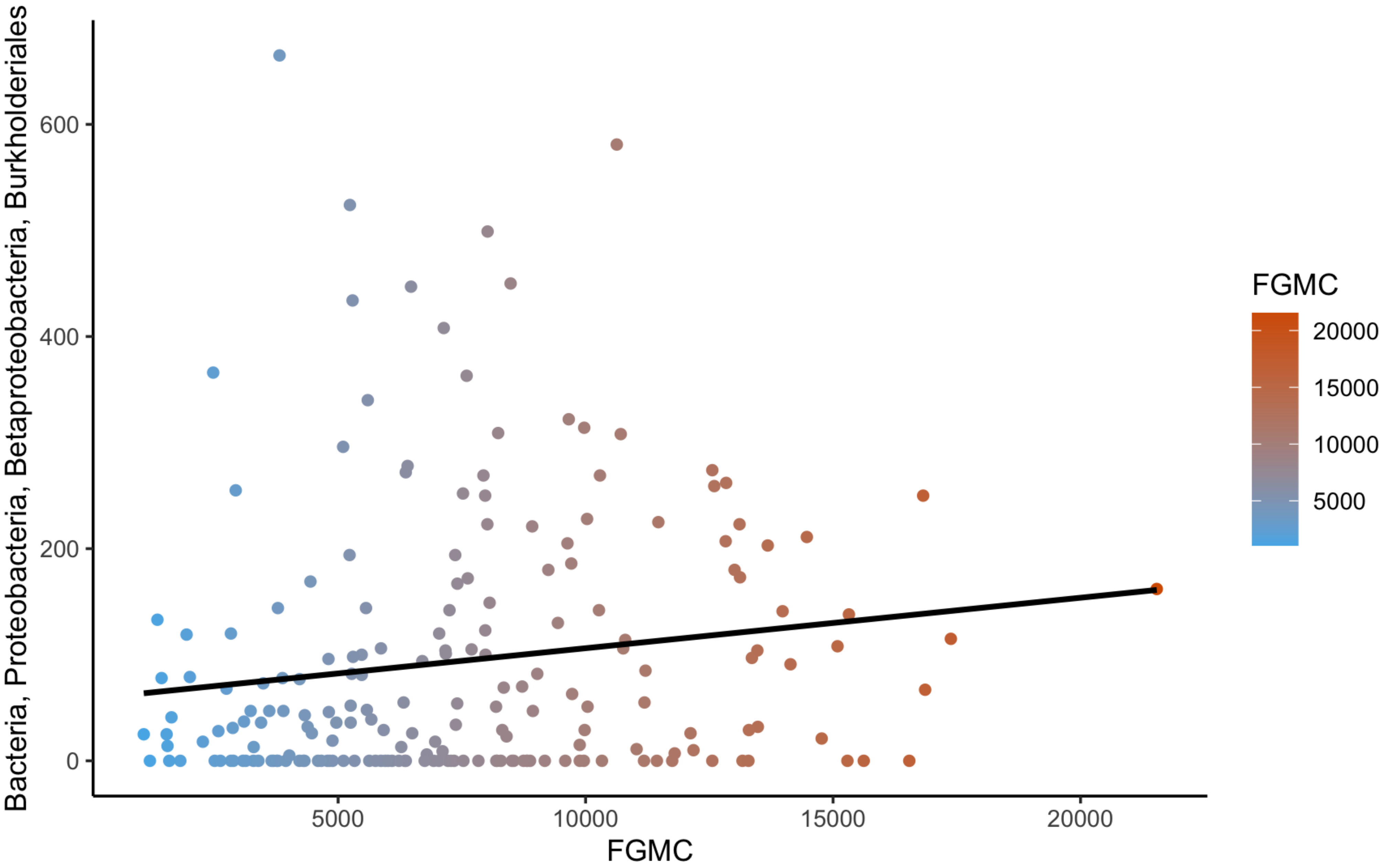

### Supplementary figure 8

**Supplementary Figure 8:**

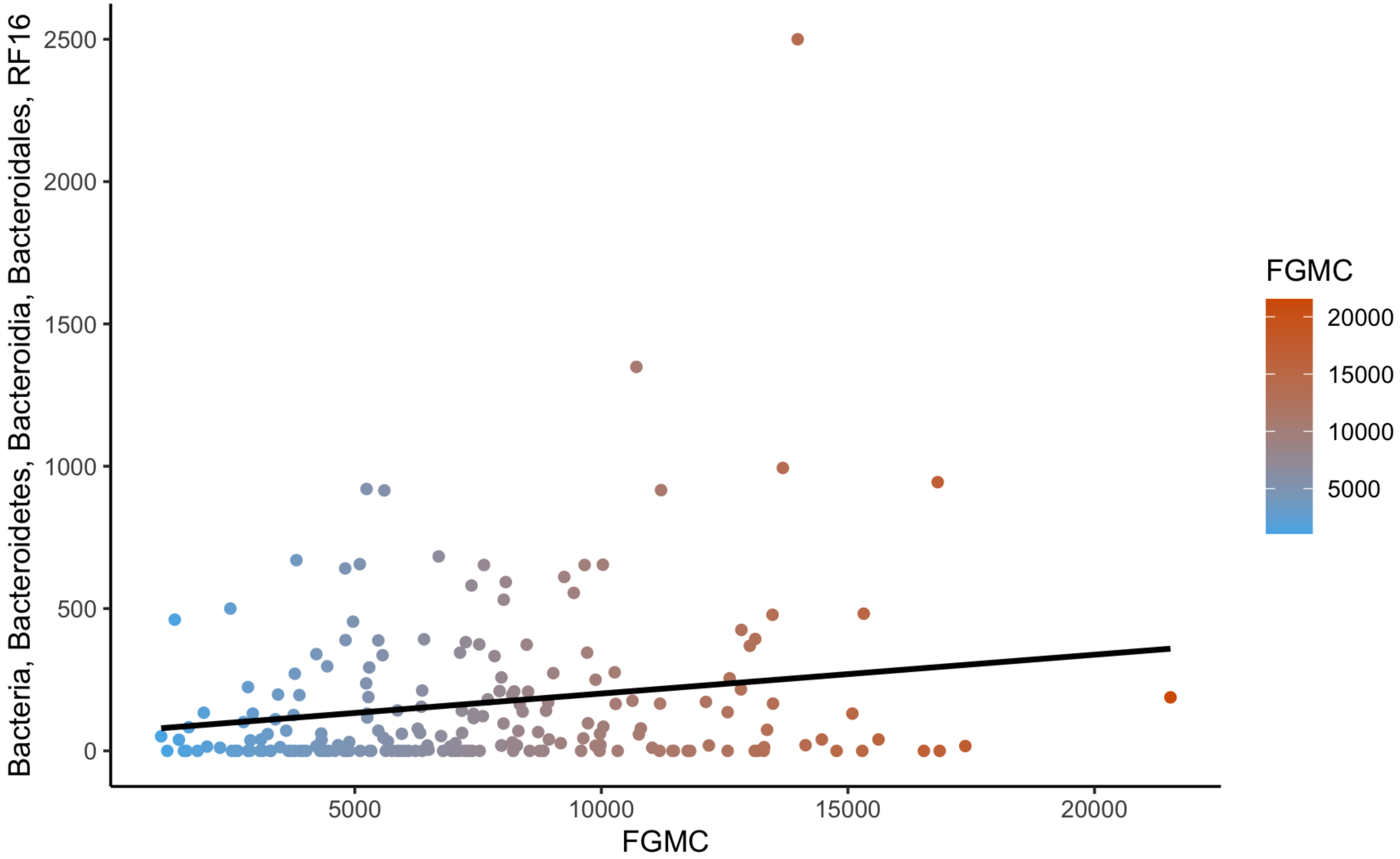

### Supplementary figure 9

**Supplementary Figure 9:**

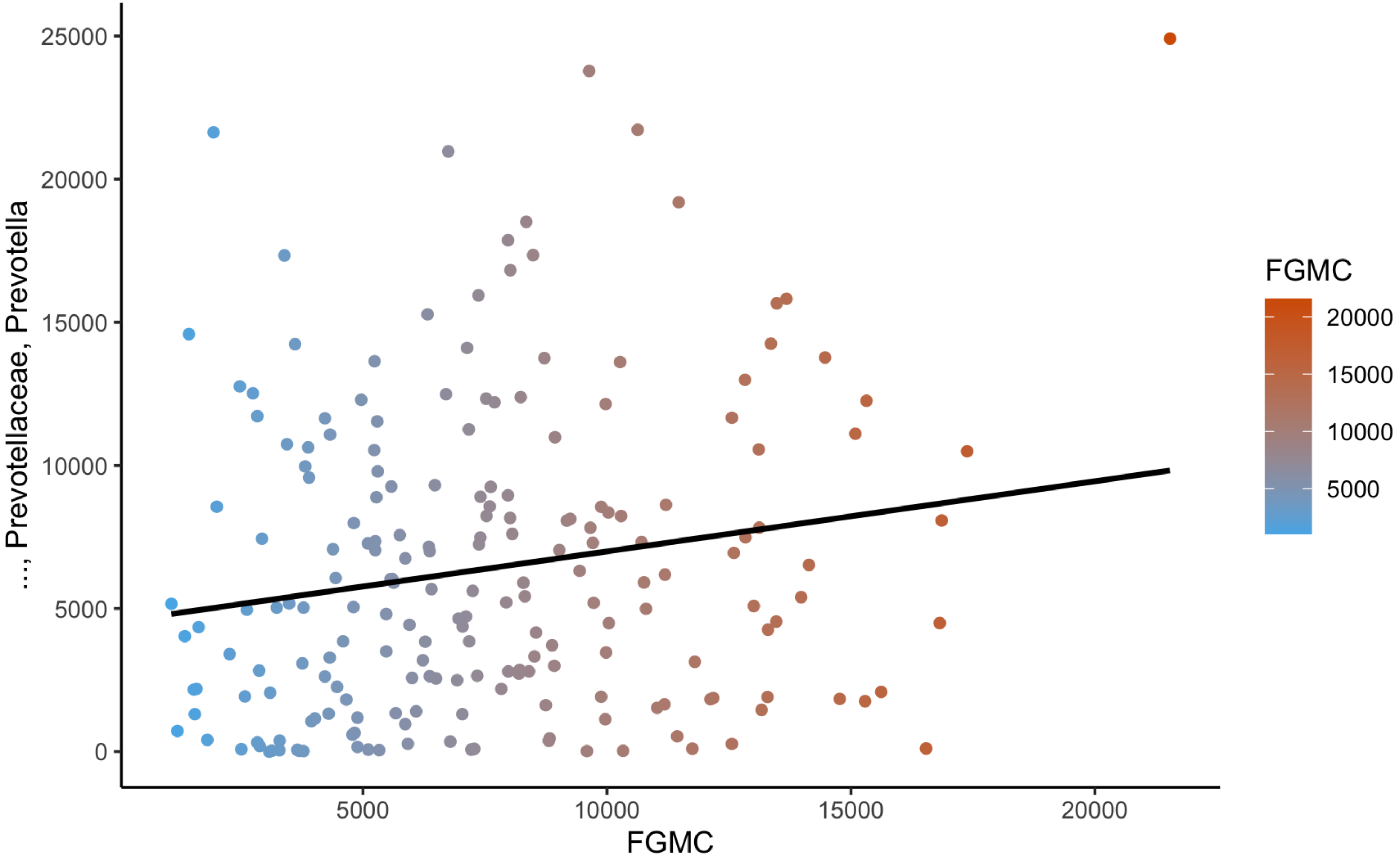

### Supplementary figure 10

**Supplementary Figure 10:**

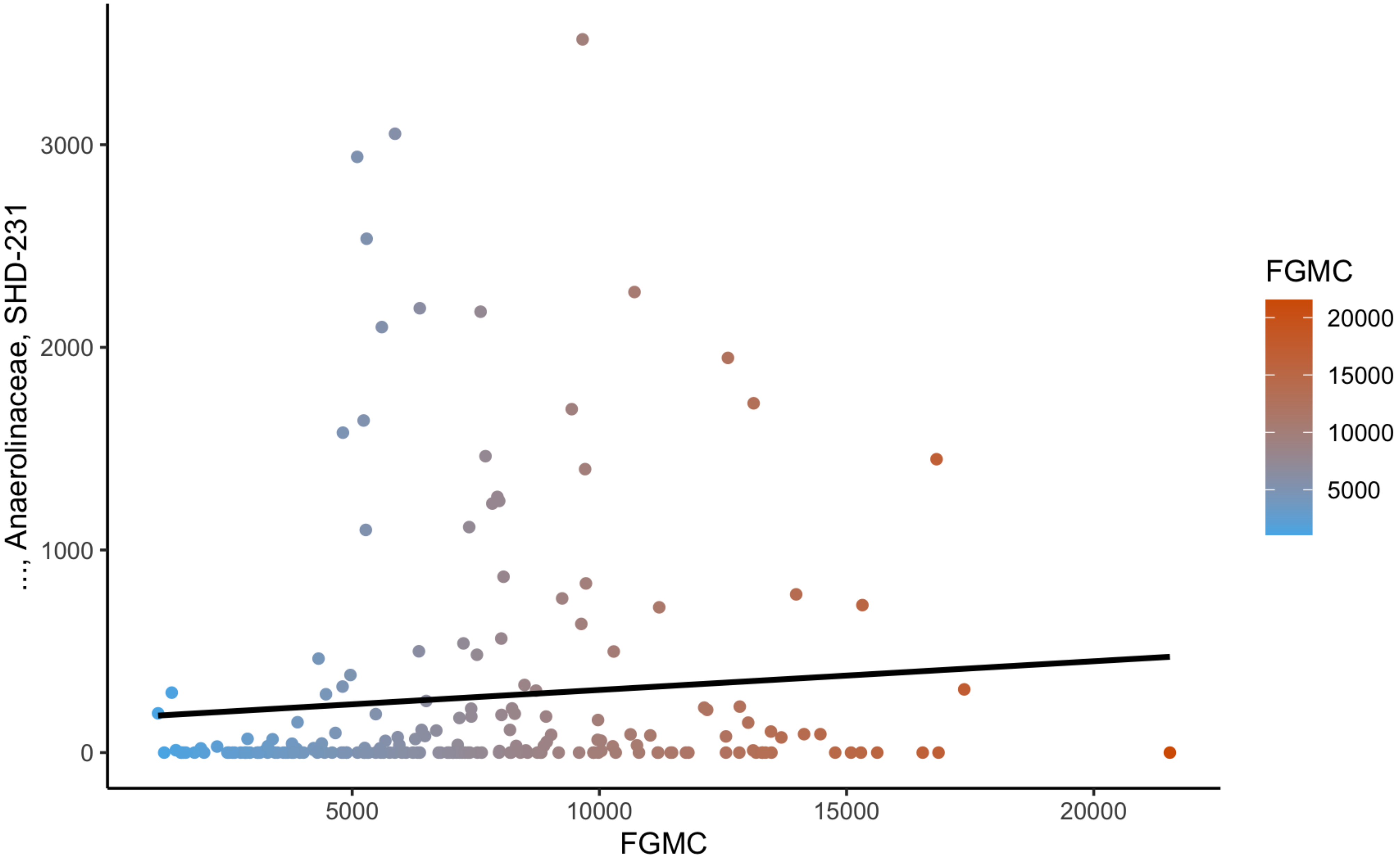

### Supplementary figure 11

**Supplementary Figure 11:**

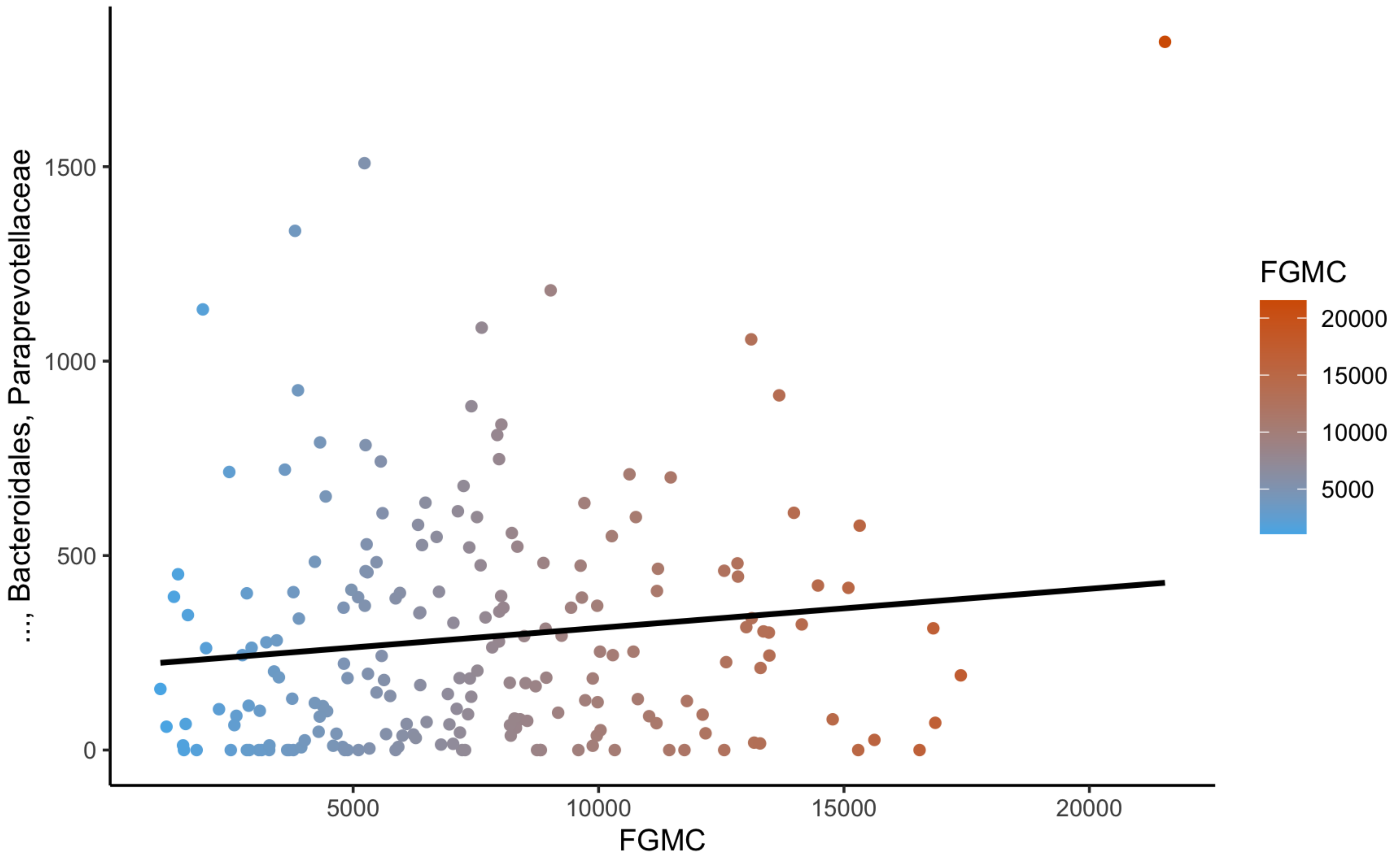

### Supplementary figure 12

**Supplementary Figure 12:**

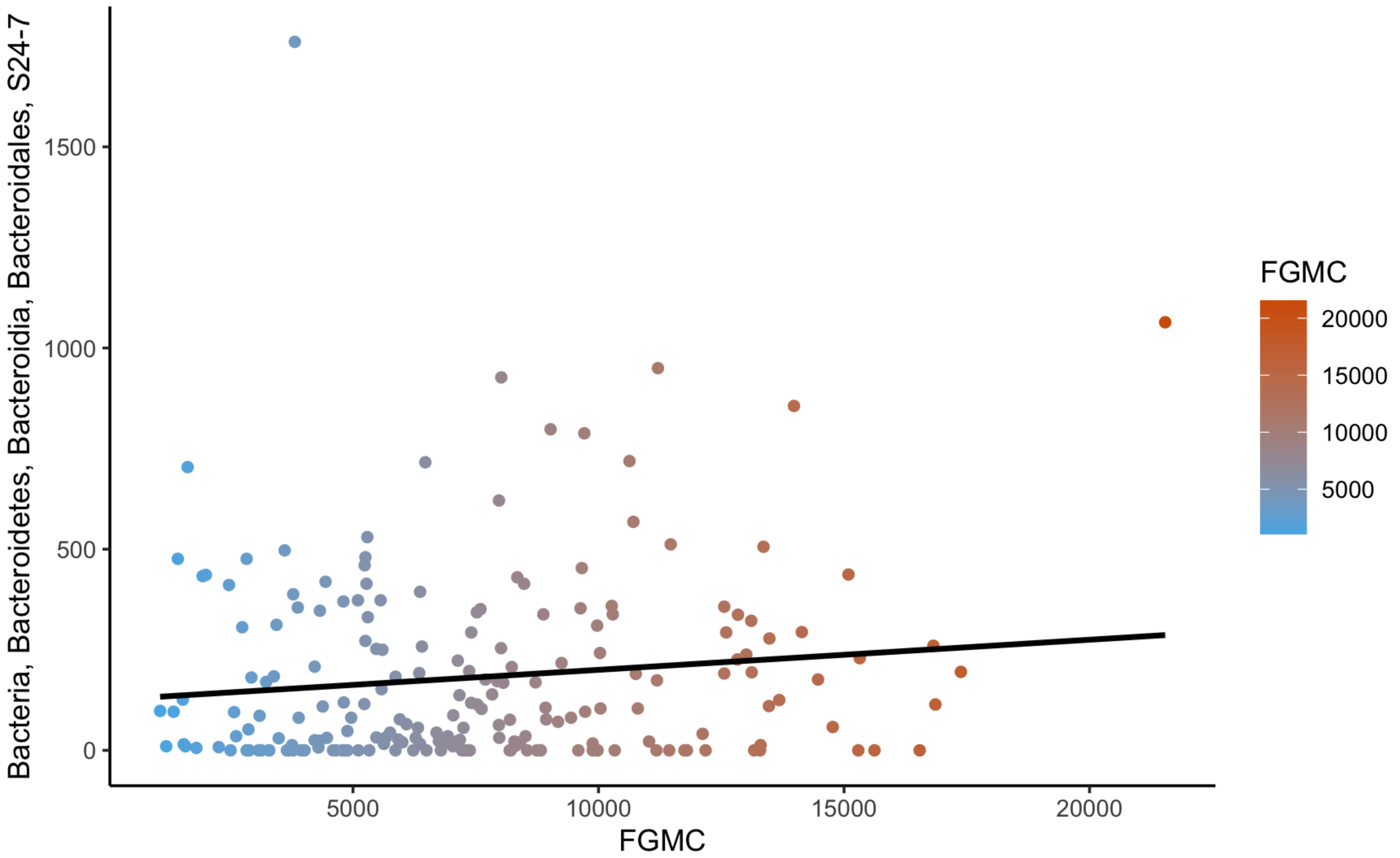

### Supplementary figure 13

**Supplementary Figure 13:**

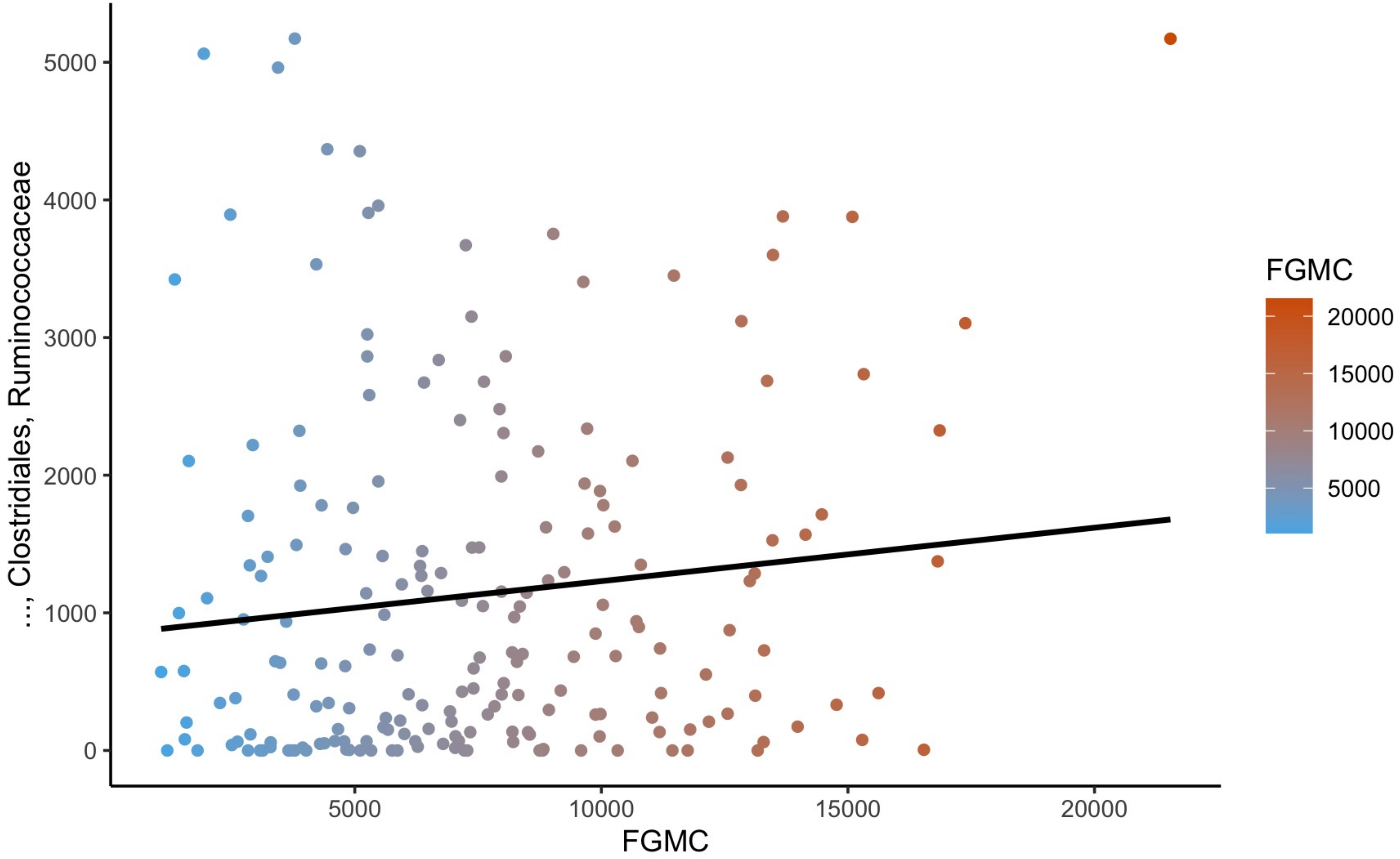

### Supplementary figure 14

**Supplementary Figure 14:**

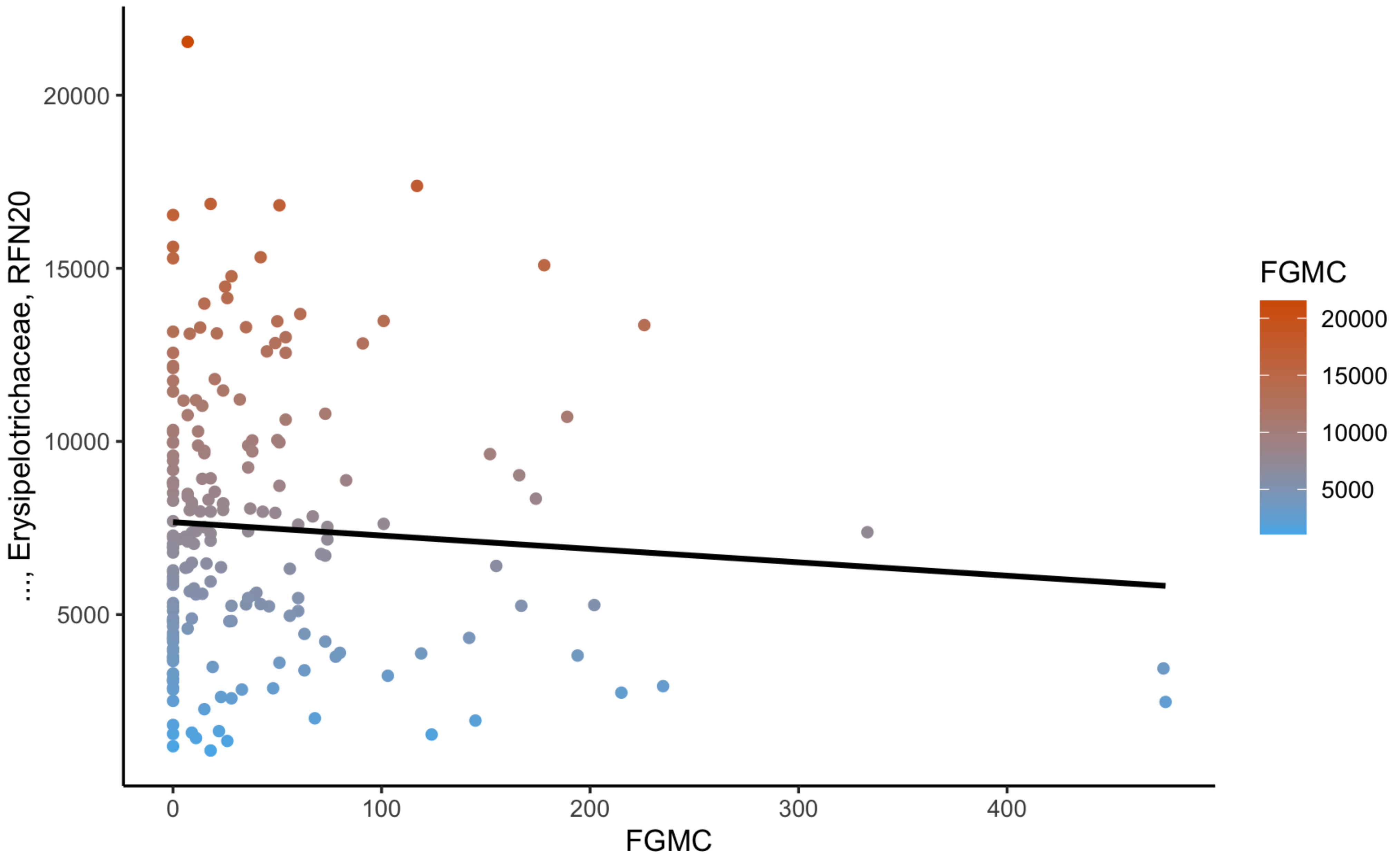

### Supplementary figure 15

**Supplementary Figure 15:**

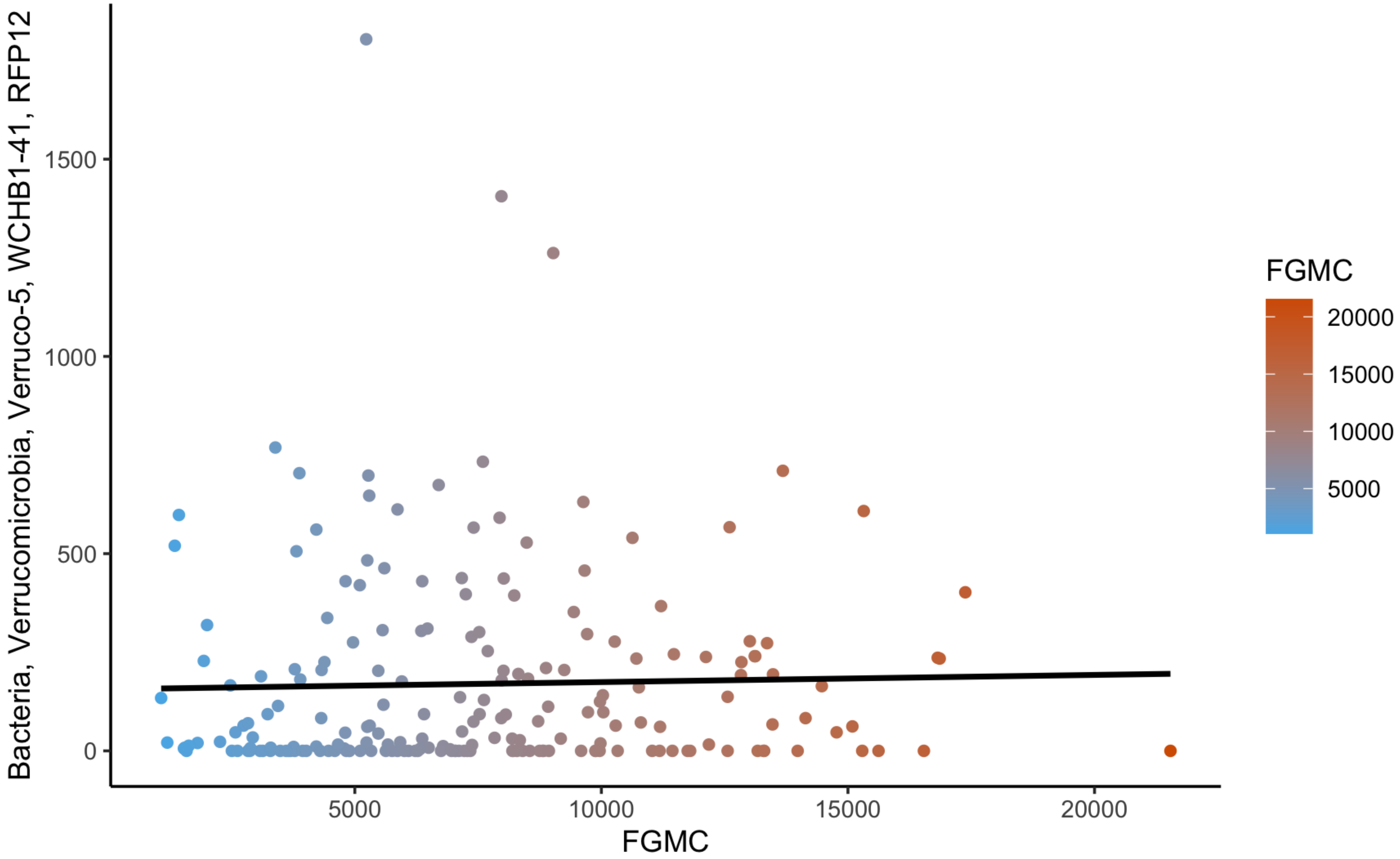

### Supplementary figure 16

**Supplementary Figure 16:**

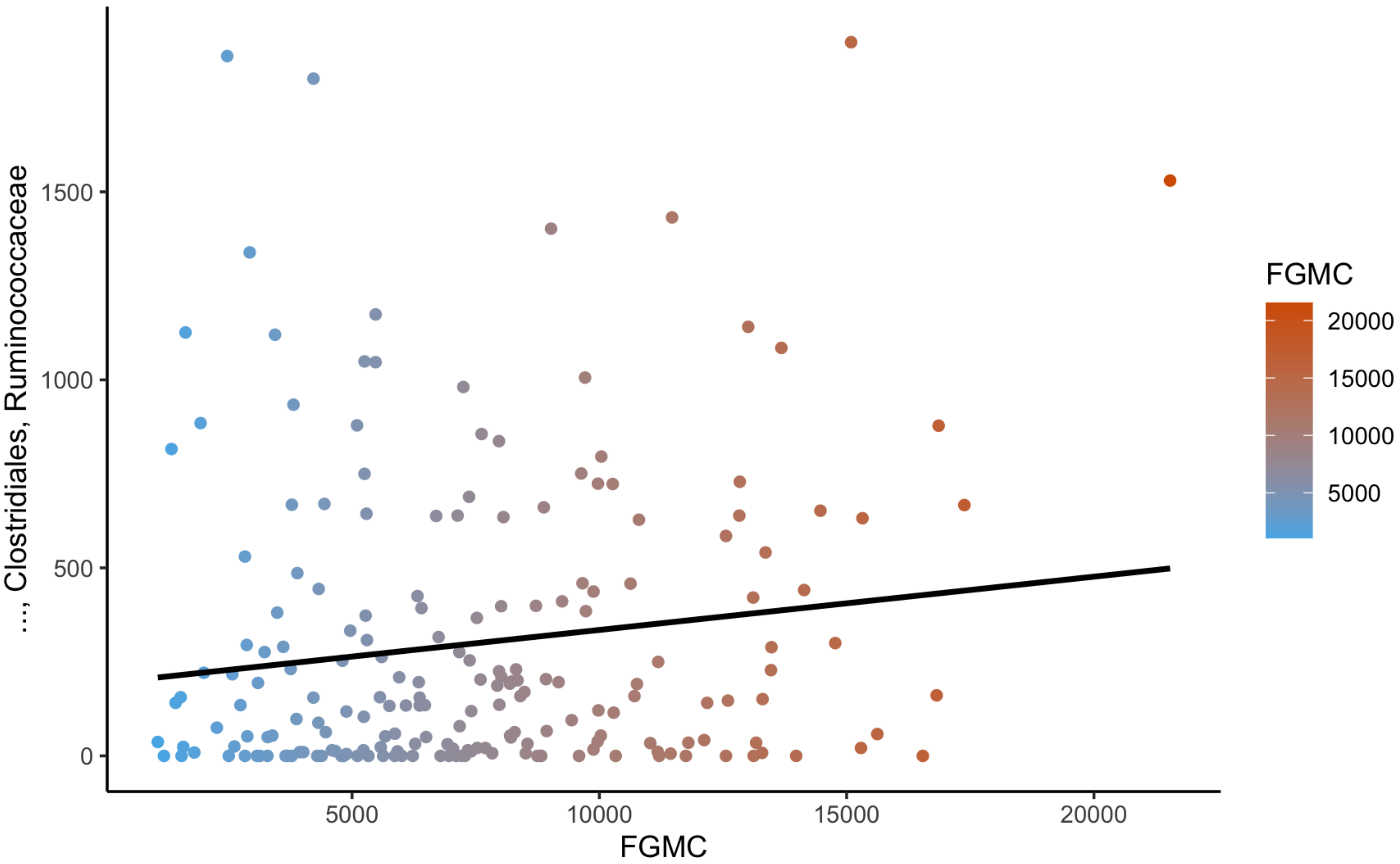

### Supplementary figure 17

**Supplementary Figure 17:**

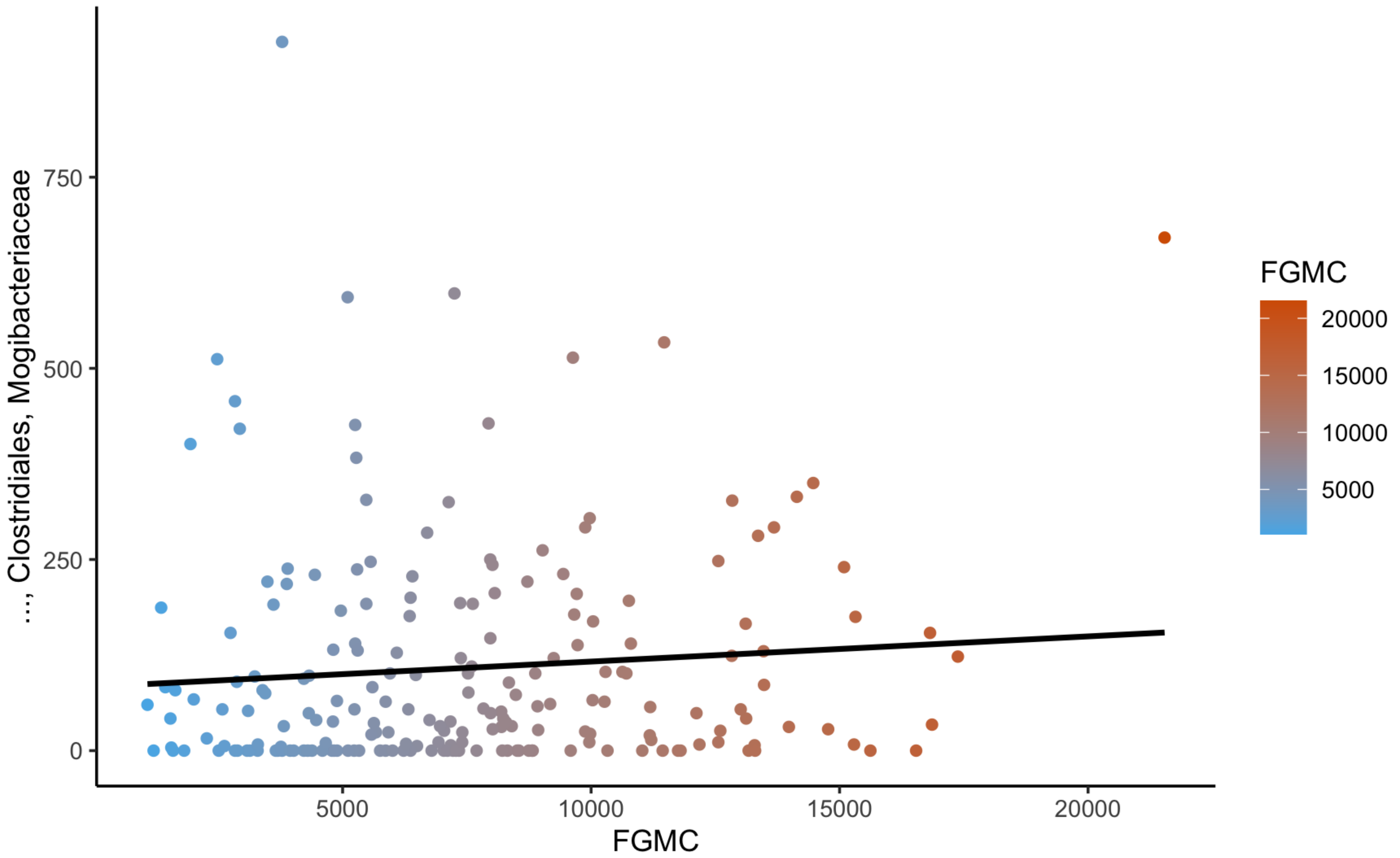

### Supplementary figure 18

**Supplementary Figure 18:**

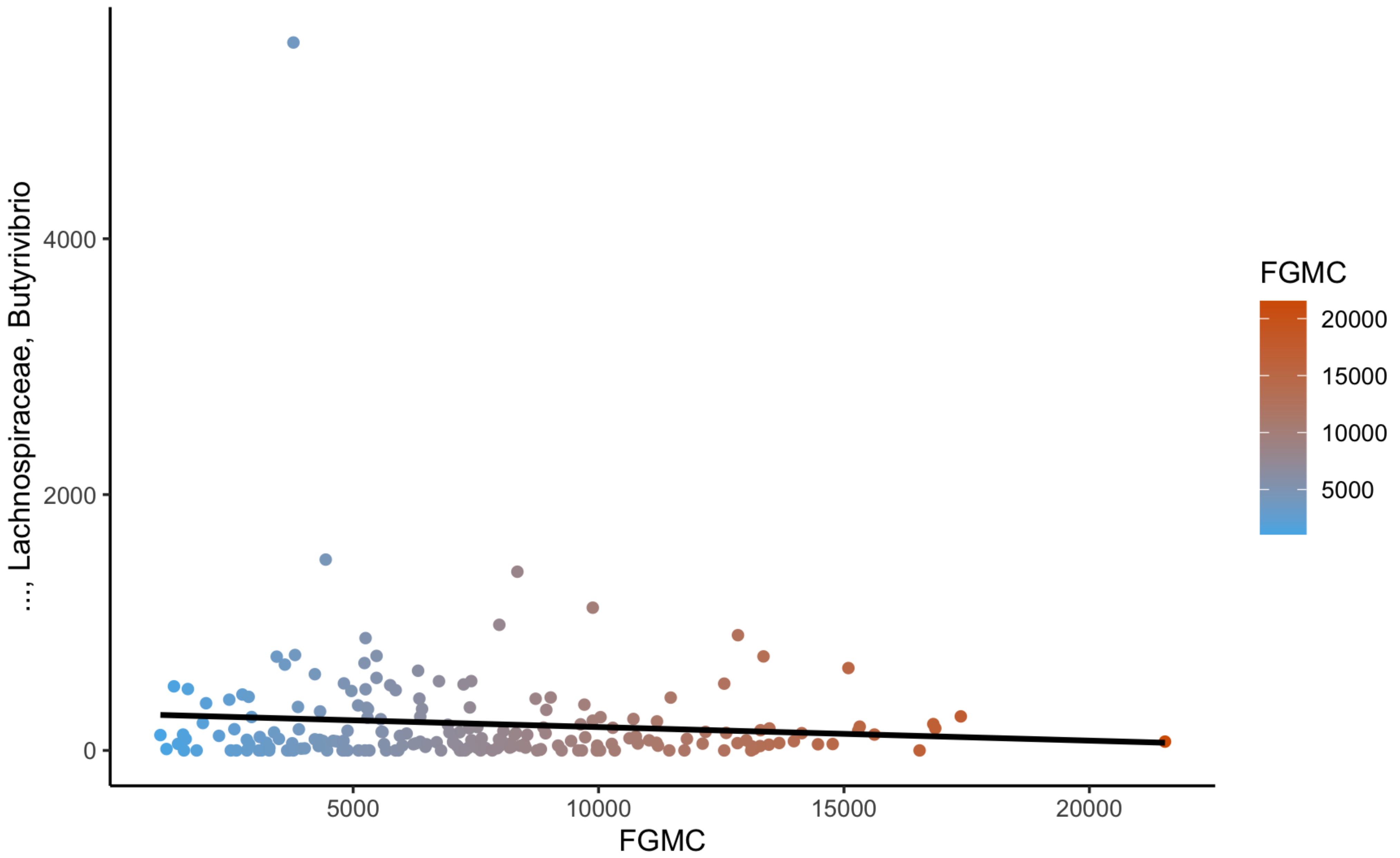

### Supplementary figure 19

**Supplementary Figure 19:**

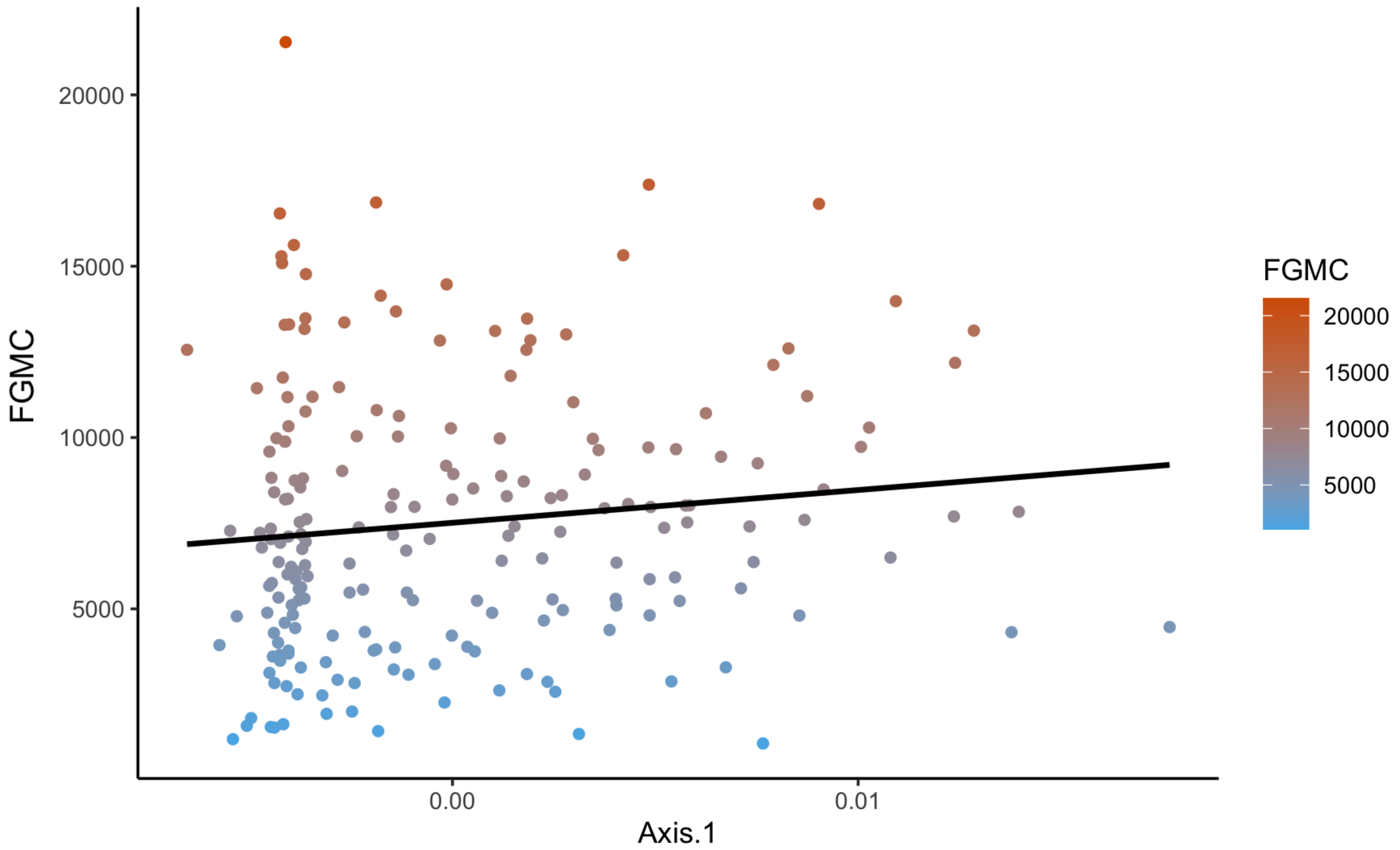
